# Directed information exchange between cortical layers in macaque V1 and V4 and its modulation by selective attention

**DOI:** 10.1101/2020.06.09.142190

**Authors:** Demetrio Ferro, Jochem van Kempen, Michael Boyd, Stefano Panzeri, Alexander Thiele

**Affiliations:** Neural Computation Laboratory, Istituto Italiano di Tecnologia, Rovereto, Italy; Center for Mind and Brain Sciences (CIMeC), University of Trento, Rovereto, Italy; Biosciences Institute, Newcastle University, NE1 7RU, Newcastle upon Tyne, United Kingdom

**Keywords:** Feedforward processing, feedback processing, visual cortex, Granger causality, attention, cortical laminae

## Abstract

Achieving behavioral goals requires integration of sensory and cognitive information, across cortical laminae and cortical regions. How this computation is performed remains unknown. Using local field potential recordings and spectrally resolved conditional Granger causality (cGC) analysis, we mapped visual information flow, and its attentional modulation, between cortical layers within and between macaque areas V1 and V4. Stimulus induced inter-laminar information flow within V1 dominated upwardly, channeling information towards supragranular cortico-cortical output layers. Within V4, information flow dominated from granular to supragranular layers, but interactions between supragranular and infragranular layers dominated downwardly. Low-frequency across-area communication was stronger from V4 to V1, with little layer specificity. Gamma-band communication was stronger in the feedforward V1 to V4 direction. Attention to the receptive field of V1 decreased communication between all V1 layers, except for granular to supragranular layers interactions. Communication within V4, and from V1 to V4, increased with attention across all frequencies. While communication from V4 to V1 was stronger in lower frequency bands (4-25 Hz), attention modulated cGCs from V4 to V1 across all investigated frequencies. Our data show that top down cognitive processes result in reduced communication within cortical areas, increased feedforward communication across all frequency bands and increased gamma band feedback communication.

## Introduction

Goal-directed behavior requires the brain to integrate sensory information with cognitive variables. In neocortical areas sensory information is conveyed by feedforward connections, while feedback connections convey information about cognitive states and goals. Feedforward and feedback connections rely on separate anatomical pathways and have been proposed to map onto distinct oscillatory frequency bands. It is, however, unknown whether these signals differ across laminae, or how they are communicated between laminae within and between cortical areas.

Feedforward connections predominantly terminate in layer IV of sensory cortical areas. This information is passed on to layers II/III and further to layers V/VI, where recurrent inputs to layer II/III arise (Callaway, 1998; Callaway, 2004; Douglas *et al.*, 1989; Douglas and Martin, 2004). Cognitive variables affect sensory processing through feedback connections, which predominantly terminate in layer I and V (Rockland and Pandya, 1979), but this termination pattern varies depending on hierarchical distances between areas (Markov *et al.*, 2014). Feedforward and feedback signals have been proposed to show separate local field potential (LFP) spectral signatures. Feedforward signals have been associated with low-frequency theta (Bastos *et al.*, 2015; Spyropoulos *et al.*, 2018) and gamma band oscillatory activity, originating and dominating in supragranular layers (Bastos *et al.*, 2015; Bollimunta *et al.*, 2011; Buschman and Miller, 2007; Lakatos *et al.*, 2008; Maier *et al.*, 2010; Smith *et al.*, 2013; Spaak *et al.*, 2012; Spyropoulos *et al.*, 2018; van Kerkoerle *et al.*, 2014; Xing *et al.*, 2012). Feedback signals have been associated with lower frequency (alpha, beta band) oscillations, prominent in infragranular layers across the cortical hierarchy (Bastos *et al.*, 2015; Buffalo *et al.*, 2011; Buschman and Miller, 2007; Popov *et al.*, 2017; Smith *et al.*, 2013; van Kerkoerle *et al.*, 2014; Xing *et al.*, 2012), although attention-related feedback signals in the gamma frequency band between FEF and V4 have been reported (Gregoriou *et al.*, 2009; Gregoriou *et al.*, 2012). Alpha related feedback has been linked to active inhibition (Bagherzadeh *et al.*, 2020; Zumer *et al.*, 2014), suggesting that feedback signals, induced by attention to the receptive field, should result in reduced alpha oscillatory power. This occurs in infragranular layers in visual areas (Buffalo *et al.*, 2011), but can also be less layer specific (van Kerkoerle *et al.*, 2014). It is thus questionable whether feedback is characterizable by alpha frequencies as attention, employing feedback, shunts alpha oscillations, instead of using them. In extrastriate sensory areas, attention increases LFP power in the gamma frequency band (Bosman *et al.*, 2012; Chalk *et al.*, 2010; Fries *et al.*, 2001; Gregoriou *et al.*, 2009; Gregoriou *et al.*, 2012; Grothe *et al.*, 2012; Grothe *et al.*, 2018; Richter *et al.*, 2017; Taylor *et al.*, 2005), while in primary visual cortex attention can increase or decrease LFP power in the gamma frequency band (Bastos *et al.*, 2015; Bosman *et al.*, 2012; Buffalo *et al.*, 2010; Buffalo *et al.*, 2011; Chalk *et al.*, 2010).

Many of the above results were obtained by methods which do not provide insight how these signals differ between laminae within an area, or between laminae across different areas. Thus, it remains unclear whether layer differences in these signals between cortical areas exist, and whether they are differently affected by cognitive goals.

To understand how information within and between areas is conveyed as a function of cognitive task, we performed simultaneous laminar recordings in areas V1 and V4 using 16-contact laminar probes while macaque monkeys performed a feature based spatial attention task. We quantified communication between laminae and areas using Granger Causality.

## Results

Monkeys performed a covert, top-down, feature guided spatial attention task. On each trial attention was directed by a central colored cue to one of 3 possible locations in a pseudo-randomized manner (Figure 1A). Monkeys had to detect a stimulus change at the cued location and ignore changes at uncued locations. To investigate how spatial attention affects interactions within cortical columns and between cortical columns of areas V1 and V4, we recorded Local Field Potentials (LFP), using 16 channel laminar probes (150 μm inter-contact spacing) in 2 adult male monkeys (62 sessions in total: 34 for monkey 1, 28 for monkey 2). We inserted probes perpendicular to the cortical surface (Figure 1B). The depth of recording contacts relative to cortical layers was determined by computing the LFP current source density (CSD, Figure 1C) (Nicholson, 1973; Nicholson and Freeman, 1975) and the multi-unit response latency (Roelfsema et al., 2007). The earliest current sink of the CSD and the shortest response latency identified input layer IV (Figure 1C). Recording sites superficial to the input layer contacts were defined as supragranular layers (L I/II/III), deeper sites were defined as infragranular layers (L V/VI) (for exact assignments see Methods) (Bollimunta *et al.*, 2008; Lakatos *et al.*, 2008; Self *et al.*, 2013; Nandy *et al.*, 2017; van Kerkoerle *et al.*, 2014). Data were analyzed for sessions in which V1 and V4 receptive fields (RF) overlapped (supplemental information for details), although centre-to-centre RF positioning could be offset (Figure 1D).

**Figure 1:**
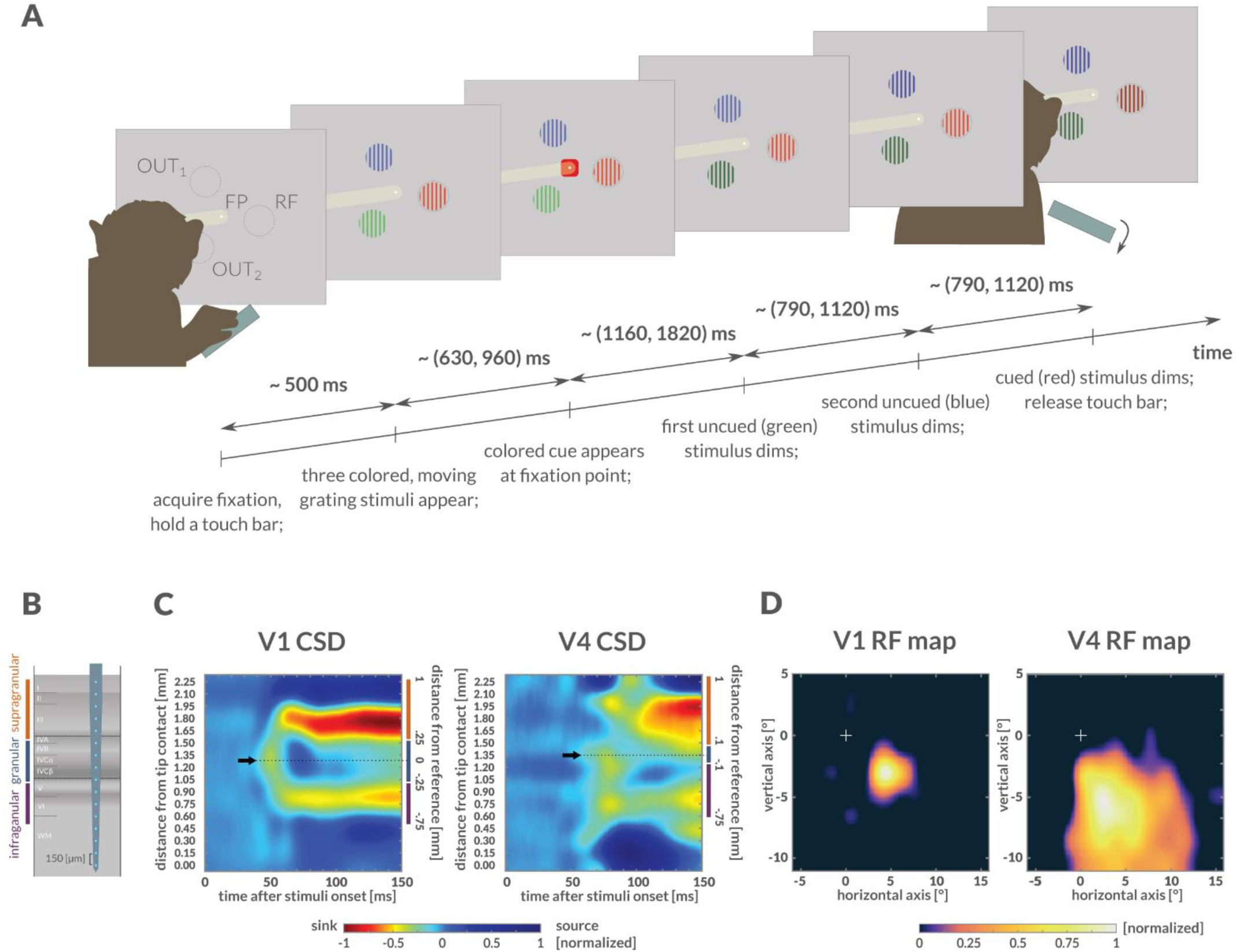
Behavioral task and recording setup. **A)** Covert, feature-guided visuospatial attention task. Monkeys fixated a fixation point (FP), and held a touch bar. Following fixation, three colored, moving grating stimuli were presented equidistant to the FP: one stimulus covered receptive field (RF) locations, the other two were located outside the RF (OUT_1_ and OUT_2_). With a random delay from stimuli presentation, a colored attention directing cue was presented at FP, indicating which stimulus was relevant on the current trial. Following the cue, the stimuli sequentially dimmed at unpredictable delays. When the relevant stimulus dimmed, the monkey had to release the touch bar to receive a fluid reward. Stimuli and cue colors, as well as the order of dimming of colored stimuli, were randomized across task trials. Ranges on the time line indicate the range of random event delays. **B)** Sketch to indicate laminar recording sites. Probes (16 contacts, 150 μm contact spacing) were injected normal to the cortical surface, aiming to cover all layers. **C)** Stimulus-induced CSD (example session), for both V1 (left) and V4 (right). Earliest current sinks were identified as layer IV (black arrows). Based on their distance from reference depth, recording contacts were assigned to granular, infragranular and supragranular compartments (shown on the right of CSDs). **D)** RF maps (mean across depths, same example session as C), for V1 sites (left) and V4 sites (right).

LFP data were analyzed in different time windows. Here we mostly present data from the time window preceding the first stimulus dimming (−503.25 to 0 ms, 512 time points). This corresponds to the period when attention was most focused on the relevant stimulus, and when attentional modulation of spiking activity is most profound in this task (Supplementary Figure S8, Thiele et al., 2016; Dasilva et al., 2019). We used bipolar re-referencing to improve spatial specificity of LFP signals (Methods for details).

In line with various reports, location of spectral power peaks differed between animals (Bosman et al., 2012; Bastos et al., 2015; Rohenkohl et al., 2018). Despite this, key analyses were performed within frequency ranges widely used in the literature (Buzsáki and Draguhn, 2004; Fries, 2005; Lakatos et al., 2008; Bosman et al., 2012; van Kerkoerle et al., 2014; Bastos et al., 2015; Richter et al., 2017; Rohenkohl et al., 2018; Spyropoulos et al., 2018). We focused on theta 4-8 Hz, alpha 8-13 Hz, beta 13-25 Hz, low-gamma 25-50 Hz, and high-gamma 50-80 Hz frequency. Adjusting frequency ranges to align with key features of individual monkey spectra (e.g. spectral peak locations) yielded qualitatively similar outcomes for all results described.

### Spectral power and coherence across V1 and V4 layers

In V1, stimulus presentation increased spectral power relative to baseline (pre-stimulus) power, across cortical layers at beta band frequencies and above (p<0.001 for beta and gamma bands for monkey 1, n=224 pooled contacts; p<0.001 for all frequency bands for monkey 2, n=257; two-sided Wilcoxon signed rank test; Figure 2A shows data pooled across layers, supplementary Figures S1, S3 show layer resolved results). Attending to the RF increased low gamma frequency peak power in monkey 1 across all layers when compared to attend out conditions (Figure 2A, supplementary Figure S1). An increase in low gamma frequency peak power was not seen in monkey 2 (Figure 2A, supplementary Figure S1). However, in both monkeys attending to the RF stimulus resulted in 3-4 Hz higher low gamma power peak location compared to attend away conditions (changes were 32.82 ± 0.30 (S.E.M) Hz to 35.58 ± 0.26 (S.E.M) Hz in monkey 1, 46.83 ± 0.15 (S.E.M) Hz to 50.63 ± 0.15 (S.E.M) Hz in monkey 2; p<0.001 both monkeys; n=224 for monkey 1, n=257 for monkey 2; two-sided Wilcoxon signed rank test; Figure 2A). This phenomenon has been described as a shift towards higher frequencies with attention (Bosman *et al.*, 2012), but it is better described as a drop in frequencies when attention is directed away from the receptive field, as stimulus presentation results in a gamma peak slightly higher than that induced by attention (Figure 2A, dashed lines). Due to the differences in peak location, attention to the RF resulted in significantly higher spectral power at frequencies above the average of attend RF and attend out peak frequency location (p<0.001 for monkey 1, n=224; p<0.001 for monkey 2, n=257; two-sided Wilcoxon signed rank tests) and significantly lower power below the average frequency (p<0.001 in beta band for monkey 1, n=224; p<0.001 in low gamma band for monkey 2, n=257; two-sided Wilcoxon signed rank tests; Figures 2A). Additionally, decreases in V1 LFP spectral power with attention were found at lower frequencies (p<0.001 for theta and alpha bands in monkey 1, n=224; p<0.05 in alpha band in monkey 2, n=257; two-sided Wilcoxon signed rank test; Figure 2A). These attentional effects were similar across cortical layers in V1 (supplementary Figures S1, S3).

**Figure 2:**
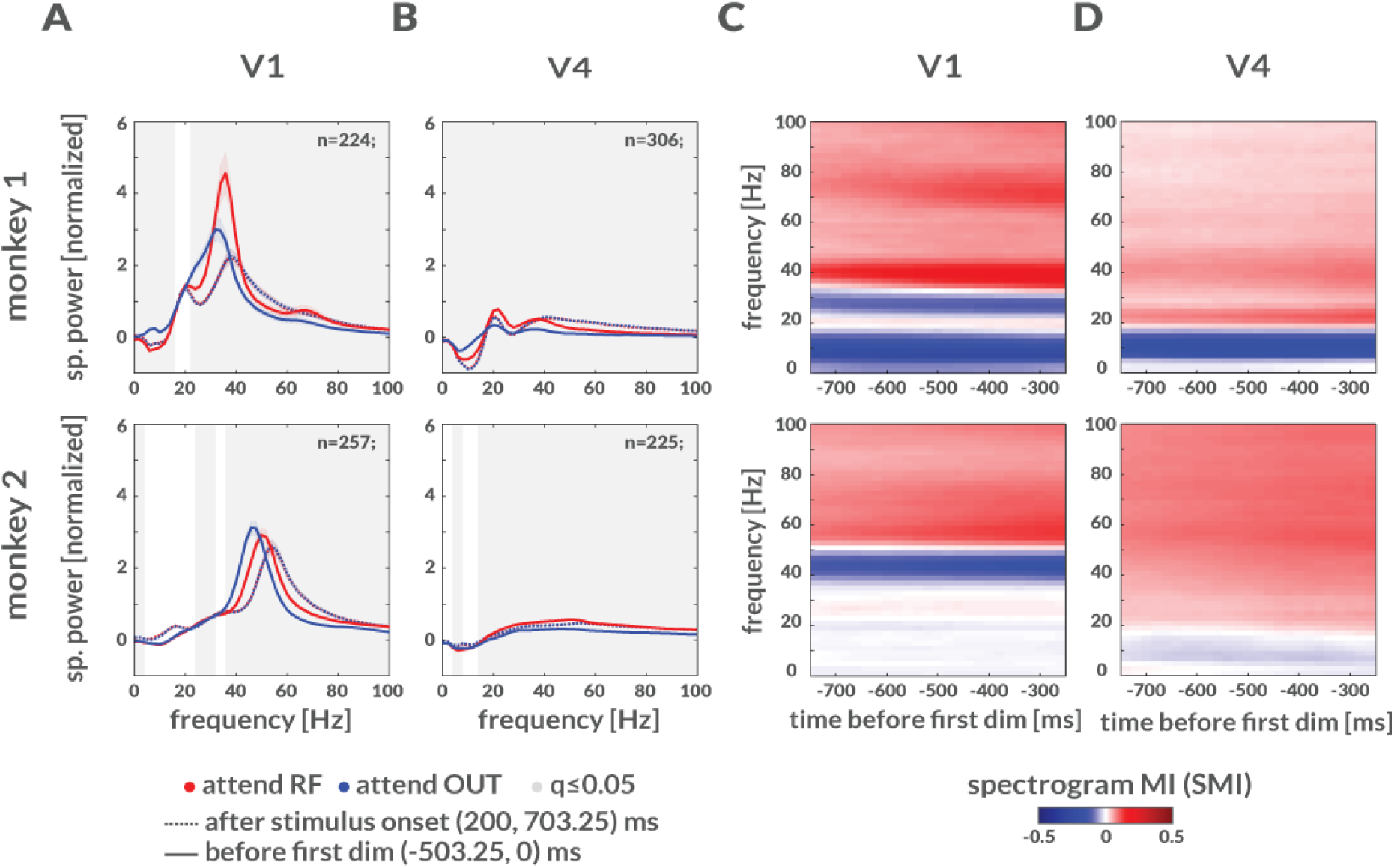
Attention decreases spectral power at lower frequencies and increases power at higher frequencies. **A)** Spectral power (mean ± S.E.M across sessions and depths) of bipolar LFP signals in ≈500 ms time windows. Dashed lines show spectral power after stimulus onset (200, 703.25) ms; solid lines show spectral power at times (−503.25, 0) ms before first dimming; shaded areas show S.E.M. Frequencies with significant difference between attentional conditions are shown by gray background (two-sided Wilcoxon signed-rank tests, FDR corrected q≤0.05). **B)** Same as in A, but for V4 LFPs. **C)** LFP attention spectral power modulation index (SMI, mean across sessions and depths) for LFPs from monkey 1 (top) and monkey 2 (bottom). Spectral analysis was applied to 503.25 ms time windows sliding in 20 ms steps, at times ≈(−1000, 0) ms before the first dimming. **D)** Same as in C, but for V4.

Stimulus onset reduced low frequency spectral power in V4 in monkey 1 but increased it in monkey 2 (<13 Hz, p<0.001 in theta and alpha bands relative to pre-stimulus power; two-sided Wilcoxon signed rank tests; supplementary Figures S2 and S3). However, in both monkeys it increased spectral power for higher frequencies (>13 Hz, beta and gamma bands; p<0.01 in monkey 1 beta band, p<0.001 in monkey 1 low and high gamma band, n=306; p<0.001 in beta and gamma bands for monkey 2, n=225; two-sided Wilcoxon signed rank tests). Attention to the RF stimulus resulted in significant increases in LFP spectral power in intermediate and high frequencies (from beta to gamma band; p<0.001 in both monkeys, n=306 contacts in monkey 1, n=225 contacts in monkey 2; two-sided Wilcoxon signed rank tests), and significant decreases at low frequencies (p<0.001 in theta and alpha bands for monkey 1, n=306; p<0.001 theta band for monkey 2, n=225; two-sided Wilcoxon signed rank tests; Figures 2B, supplementary Figures S2, S3). In V4, effects of attention on spectral power were largely similar across cortical layers in both monkeys (supplementary Figures S2, S3).

To assess attentional modulation of spectral power relative to the time of cue onset and to the time of the first dimming we calculated spectrogram modulation indices (SMIs) using a sliding window of 512 time points (503.25 ms length, Methods). Attentional modulation of spectral power (either positive or negative) increased after cue onset and persisted until the time of first dimming (supplementary Figures S1-S3). In V1, SMIs were positive for higher gamma frequencies, showed negative SMI for a narrow frequency just below the average gamma peak, followed by positive SMIs in the beta band and negative SMIs in low frequency ranges (alpha and theta band, Figure 2C). In V4, SMIs were negative for low frequency spectral power, i.e. attention reduced low frequency power in V4, while they were positive for frequencies >15-20 Hz, i.e. attention increased spectral power for mid and high frequencies (Figure 2D).

Attentional modulation of intra-area LFP spectral coherence largely followed the pattern described for spectral power (Figures 3A-B). This indicates that the local (bi-polar referenced) LFP power at specific frequencies is tightly coupled between layers. Attention to the RF resulted in significantly ≈1-2 Hz higher spectral coherence peak locations in the gamma-band in V1 (from 35.53 ± 0.13 (S.E.M) Hz to 36.50 ± 0.12 (S.E.M) Hz in monkey 1, from 47.53 ± 0.06 (S.E.M) Hz to 49.61 ± 0.06 (S.E.M) Hz in monkey 2; p<0.001 in both monkeys, n=1100 contact pairs for monkey 1, n=1512 for monkey 2; two-sided Wilcoxon signed rank tests; Figure 3A), it increased spectral coherence at higher frequencies (p<0.001 in low and high gamma in monkey 1, n=1100 contact pairs; p<0.001 high gamma in monkey 2, n=1512; two-sided Wilcoxon signed rank tests; Figure 3A) and decreased coherence at lower frequencies (p<0.001 in theta and alpha bands, p<0.05 in beta band for monkey 1, n=1100; p<0.001 in beta and low gamma bands in monkey 2, n=1512; two-sided Wilcoxon signed rank tests; Figure 3A). Slight increases were also found in lower bands (p<0.05 in lower beta band within ≈16-18 Hz in monkey 1, n=1100; p<0.001 in theta band for monkey 2, n=1512; Figure 3A). In V4 spectral coherence was increased by attention at higher frequencies (beta and gamma bands, p<0.001 in both monkeys; n=1949 contact pairs in monkey 1, n=1295 in monkey 2; two-sided Wilcoxon signed rank tests; Figure 3B), and decreased at lower frequencies (p<0.001 in theta and alpha bands in monkey 1, n=1949; p<0.01 in theta band in monkey 2, n=1295; two-sided Wilcoxon signed rank tests; Figure 3B). Inter-areal spectral coherence showed three main peaks (Figure 3C). One peak occurred at low frequencies (theta/alpha band), where attentional modulation differed between monkeys for the theta, but not for the alpha band (coherence was decreased in theta band for monkey 1, p<0.001, n=1940; increased in alpha band for monkey 1, p<0.001, n=1940; increased in theta band p<0.05, and alpha band p<0.001 for monkey 2, n=1802; two-sided Wilcoxon signed rank tests). A second peak occurred in the beta band, with increased coherence for attend RF conditions (p<0.001 in both monkeys; n=1940 in monkey 1, n=1802 in monkey 2; two-sided Wilcoxon signed rank tests). A third peak occurred in the gamma band which increased for attend RF conditions (p<0.001 in low gamma for both monkeys, p<0.001, n=1940 in high gamma in monkey 1; p<0.001, n=1802 in monkey 2; two-sided Wilcoxon signed rank tests; Figure 3C). The effects of attention on spectral coherence were largely similar between layer pairs within areas, as well as across layer pairs between areas (supplementary Figure S4).

**Figure 3:**
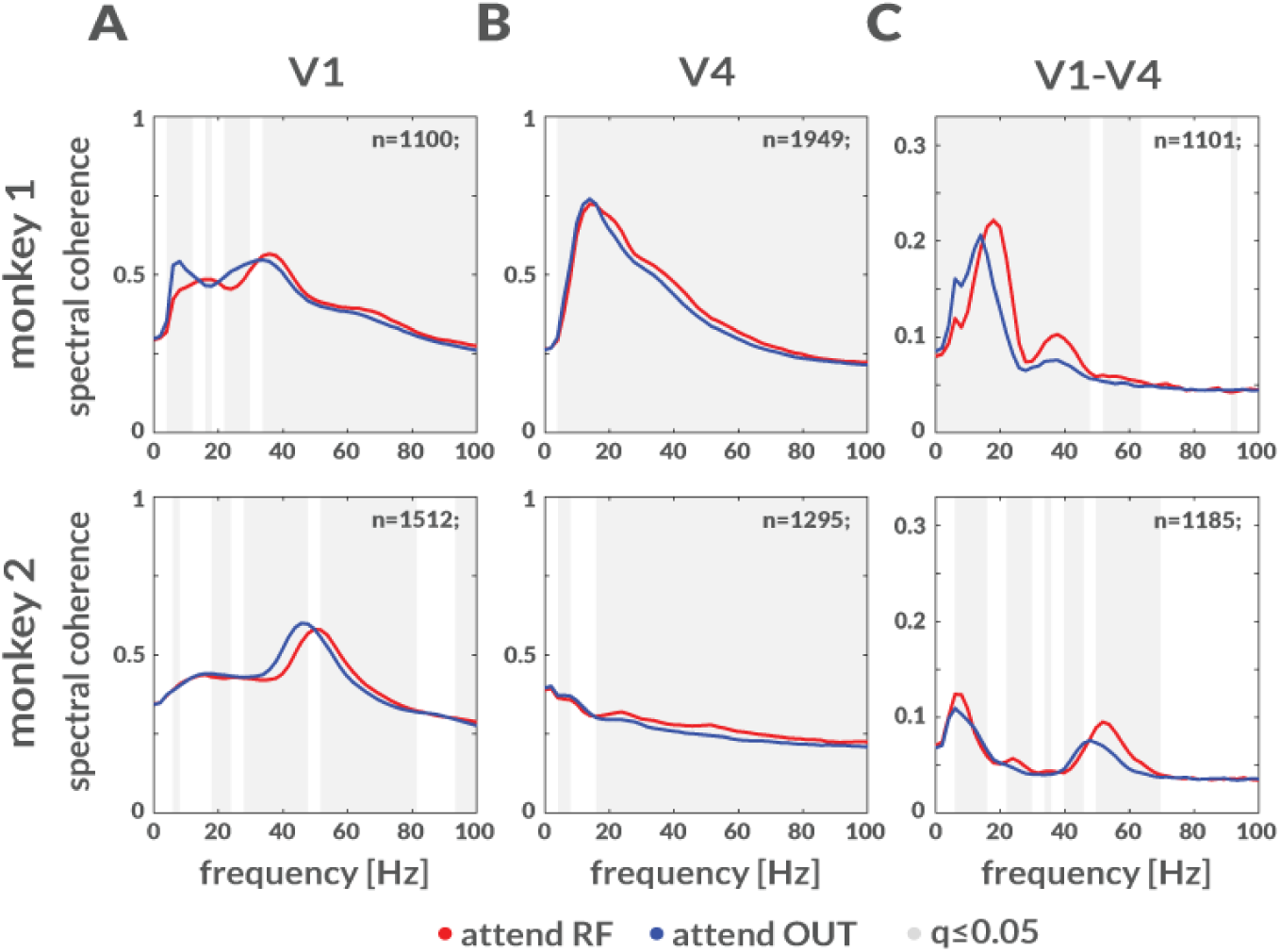
Effect of attention on LFP spectral coherence. **A) LFP** spectral coherence across V1 depths (mean ± S.E.M across sessions and depth pairs) at times (−503.25, 0) ms before the first dimming for monkey 1 (top) and monkey 2 (bottom). Gray background shows frequencies with significant difference between attentional conditions (two-sided Wilcoxon signed-rank tests, FDR corrected q≤0.05). **B)** Same as in A, but for V4. **C)** Same as in A, but for V1-V4 coherence.

### Causal communication between cortical layers and between cortical areas

To determine the flow of information within and between layers within and between areas we calculated conditional Granger Causality (cGC). cGC (Geweke, 1984) is a multivariate directed measure that allows to quantify the degree of causal relationship (or communication) between two nodes. For any directed contact pair (X,Y), cGC yields a conditional estimate of the causal flow from Y to X (and from X to Y), with the aim to discount the indirect influence of time-lagged interactions with contacts not covering the same laminar compartments as X and Y (Methods).

We first describe dominant interactions between layers and areas, irrespective of the effects of attention. This provides insight which frequency bands predominantly carry feedforward and which frequency bands predominantly carry feedback information, independent of changing cognitive variables. Spectrally resolved intra-areal and inter-areal cGCs averaged across contact pairs are shown in Figure 4. All cGCs were significant (significance threshold is shown by dashed line in Figures 4A-D, computed as 95th percentile of cGCs with trials randomly shuffled; Methods).

**Figure 4:**
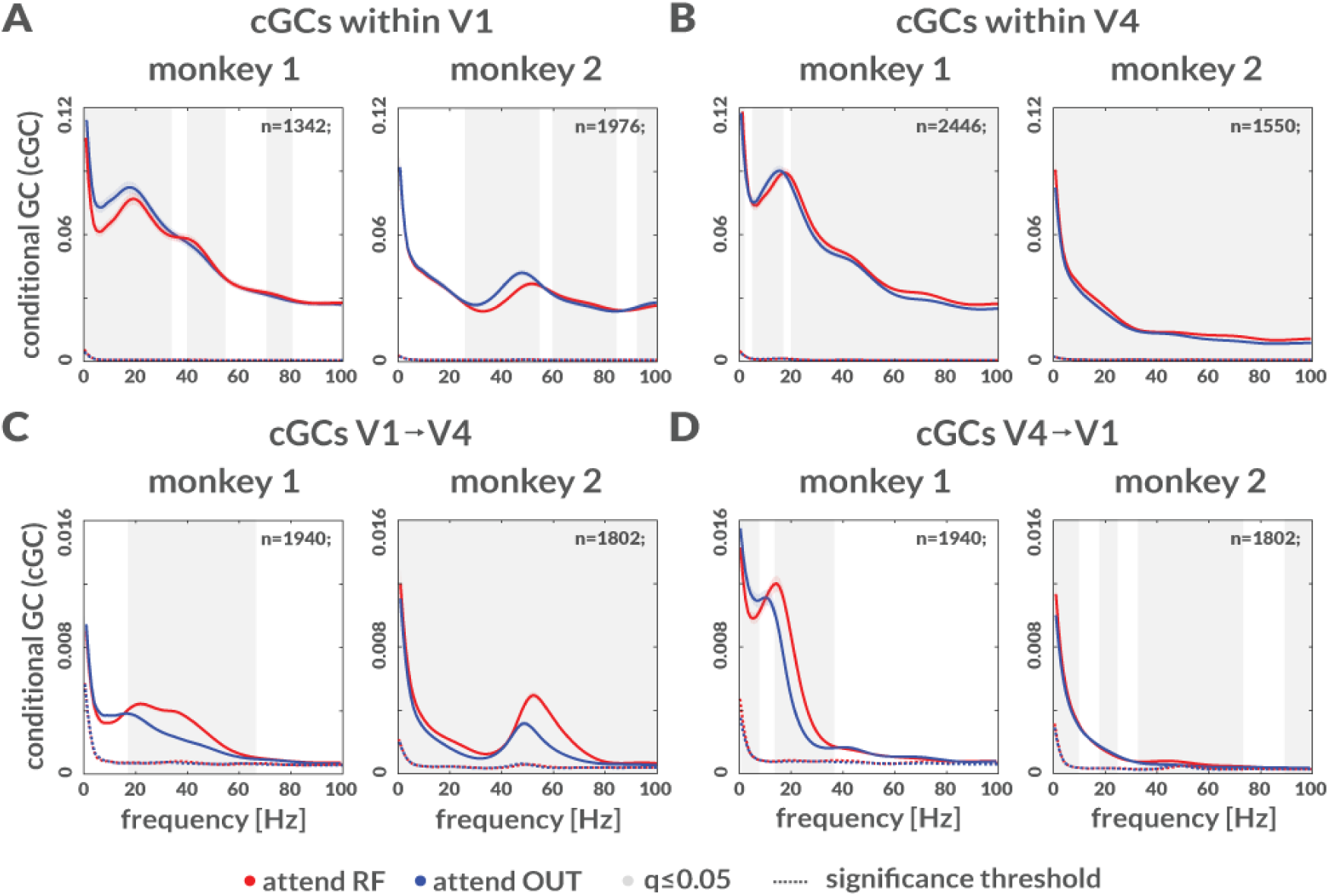
Conditional Granger Causality (cGC) for LFP signals. **A)** Conditional cGC pooled across depths in V1 (mean ± S.E.M across sessions and directed depth pairs) at times (−503.25 to 0) ms before first dimming for monkey 1 (left) and monkey 2 (right). Gray background shows frequencies with significant difference between attentional conditions (Wilcoxon signed-rank tests, FDR corrected q≤0.05,). **B)** Same as in A, but for V4. **C)** Same as in A, but for cGC from V1 to V4. **D)** Same as in A, but for cGC from V4 to V1.

To plot cGC results, we normalized each cGC to the maximum cGC across the 5 frequency bands (separately for within area and between areas cGCs after averaging across all sessions) for each monkey. To assess the dominant directionality of communication, for each contact pair (X,Y) we determined whether cGC was stronger from X to Y, or whether it was stronger from Y to X, and whether the directional difference was significant for a given frequency range (q<0.05, two-sided Wilcoxon signed rank tests, FDR corrected within frequency bands). We only present contact pairs where the directional cGC difference was significant. Significant differences are reported with color code indicating the dominant directions. For example, if a granular to supragranular cGC was larger than vice versa, it will be displayed in green in the cGC matrices, while the inverse direction will be displayed in magenta (Figure 5; for all contact cGCs differences, including non-significant ones see supplementary Figure S6). Color intensity shows the relative strength of the interactions.

**Figure 5:**
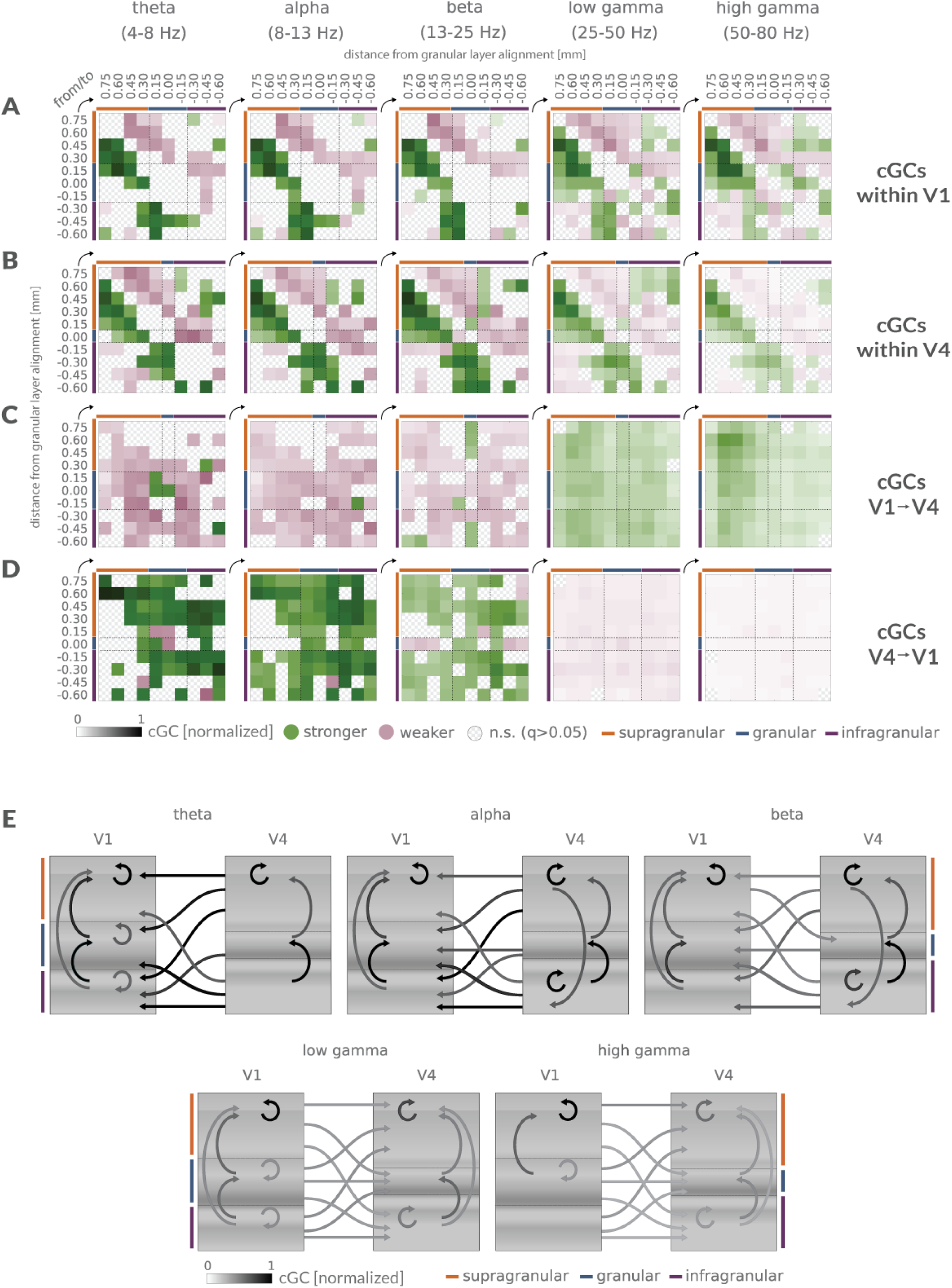
Directed connection matrices and influencer diagram of dominant cGC interactions before the first dimming. **A)** Normalized cGCs matrices (mean across sessions, pooled for the two monkeys) within V1 columns, for different frequency bands. Connection matrices are color coded to show significant dominant directions (green) and the opposite weaker directionality (magenta). Color intensity shows the relative cGC strength. Significance of cGCs difference was assessed by two-sided Wilcoxon signed rank tests, FDR corrected (*q*≤0.05) within frequency bands. cGCs were first normalized for each monkey to the peak magnitude across frequencies, then pooled. **B)** Same as in A, but cGCs for V4. **C)** Same as in A, but for V1 to V4 pairs. **D)** Same as in A, but for V4 to V1 pairs. **E)** Influencer diagram of significant dominant cGC interactions, summarizing results in A-D. Arrows show dominant cGC interactions pooled for the two monkeys, averaged for the three laminar compartments (supragranular, granular, infragranular). Gray scale intensity of arrows indicates relative strength of cGCs (independently normalized for directions within V1, within V4, and between V1 and V4).

In V1, cGCs dominate in an upwards direction within supragranular layers for all frequencies (Figure 5A), they dominate in an upwards direction for all frequencies from granular to supragranular contacts, and they dominate in an upwards direction from infragranular to granular and supragranular contacts in the theta, alpha and beta frequency range, with smaller directional differences in the gamma frequency ranges. This pattern suggests that dominant interactions converge onto feedforward cortico-cortical output (supragranular) layers.

In V4 (Figure 5B), dominant interactions occurred in an upward direction within supragranular layers, across all frequency bands. Additionally, dominant cGCs were present in an upward direction from granular to supragranular layers, and from infragranular to granular layers. However, unlike in V1, cGCs dominated in a downward direction from supragranular to infragranular layers for most contacts and frequencies. Thus, within V4, a bidirectional dominance was found, whereby directly neighboring compartments communicated more strongly in an upward direction, while more distant compartments communicated more strongly in a downward direction.

Interactions between V1 and V4 were dominated in the feedback direction in lower (theta to beta) frequency bands (magenta color dominates for these frequency bands in Figures 5C- and in the feedforward directions in the gamma frequency ranges (green color dominates for these frequency bands in figure 5C-D). These V1-V4 interactions had little layer specificity with respect to origin or destination.

cGC interactions from V4 to V1 were strongest in theta to beta frequency bands (Figure 5D). In the theta and alpha band, they were most pronounced from V4 supragranular to all V1 layers. Strong interactions also occurred from V4 infragranular to V1 infragranular layer (Figure 5D). In comparison, V4 to V1 cGCs in the gamma frequency ranges were small (even though they were significant). Thus, the feedback cGC interactions predominantly occurred in lower frequency bands, they originated in V4 supra- and infragranular layers and affected V1 supra- and infragranular layers.

These intra- and inter-areal cGC interactions are summarized in an ‘influencer’ diagram (Figure 5E). It shows that in V1 dominant communication across almost all frequencies occurs in an upwards direction towards the supragranular cortico-cortical output layer. In area V4 dominant communication occurs in a circular manner for lower frequencies (theta to beta), upwards within compartments and between neighboring compartments, but downwards from supragranular layers onto infragranular layer. In the gamma frequency range, dominant V4 communications were directed upwards towards the supragranular cortico-cortical output layer, mirroring the effects seen in V1. In the theta to beta frequency range, almost all interactions between V1 and V4 dominated in the feedback direction, while feedforward cGCs significantly dominated in the gamma frequency range.

### Attentional modulations of cGC interactions

To assess attentional modulation of cGCs, we calculated modulation indices (MIs) for each recording and determined whether MIs of cGCs between layer compartments were significant (q<0.05, two-sided Wilcoxon signed rank tests, FDR corrected within frequency bands, Methods). Figure 6 shows significant intra- and inter-areal contact pairwise cGC attentional MIs pooled for the two monkeys. Figure 7 shows significant cGC attentional MI averaged by laminar compartments (supragranular, granular, infragranular; for all contact cGCs MIs differences, including non-significant ones see supplementary Figure S7). Surprisingly, within V1 cGC MIs were mostly negative, indicating that attention reduced cGCs in an upward and a downward direction across frequency bands (Figures 6A, 7A). except from granular to supragranular contacts. Additionally, high gamma band cGCs increased with attention from granular to supragranular layers, from supragranular to granular contacts, and from granular to infragranular contacts (Figures 6A, 7A). The predominant reduction in cGC with attention within V1 was surprising given that attention increased neuronal firing rates across layers within V1 (supplementary Figure S8).

**Figure 6:**
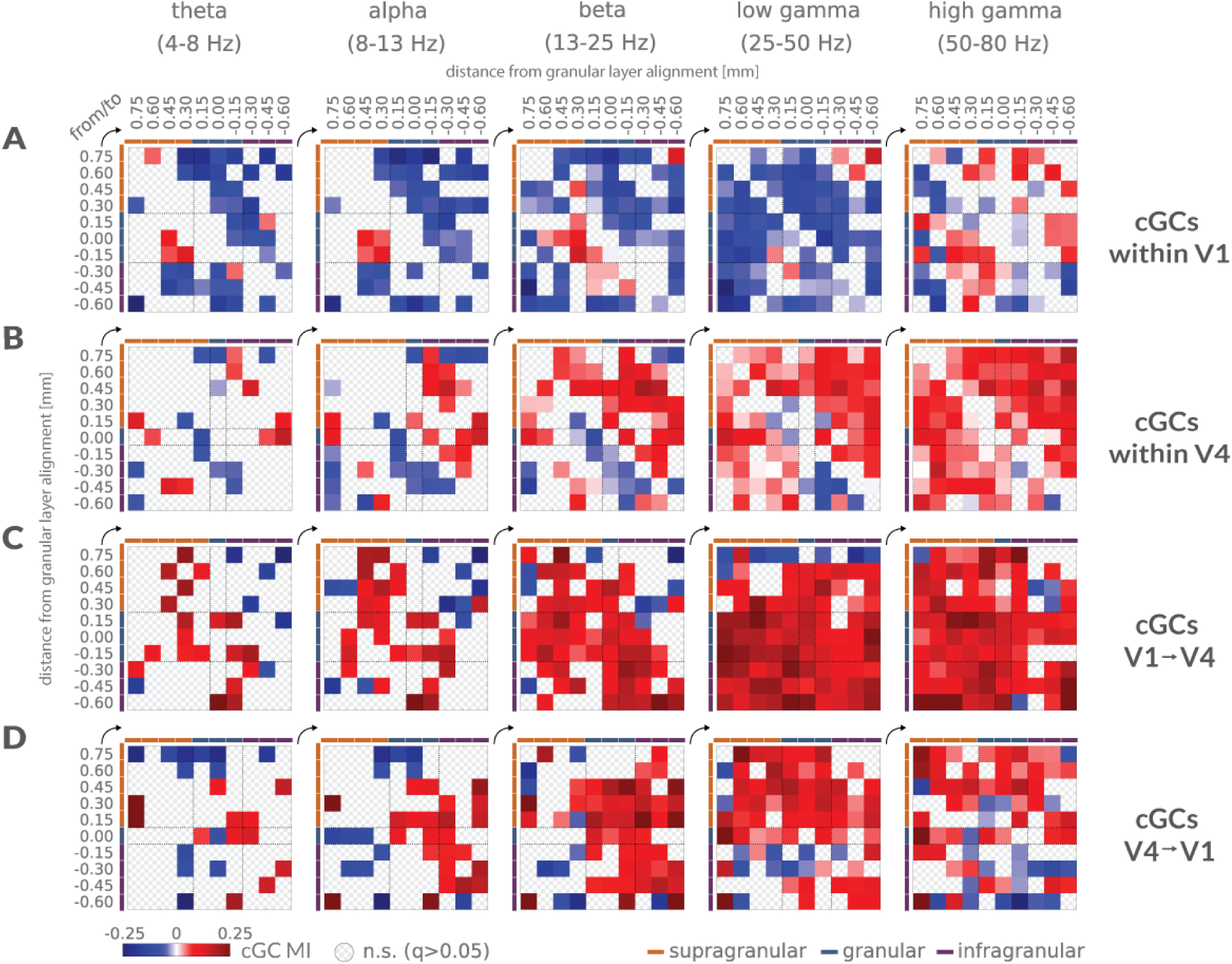
Attentional modulation of cGCs. **A)** Significant attentional modulation index of cGC (cGC MI) among depth pairs within V1 (mean across sessions, pooled for the two monkeys), at different frequency bands (significance assessed via two-sided Wilcoxon signed rank tests, FDR corrected (q≤0.05) within frequency bands). **B)** Same as in A, but for V4. **C)** Same as in A, but for cGCs MIs from V1 to V4. **D)** Same as in A, but for cGCs MIs from V4 to V1.

**Figure 7:**
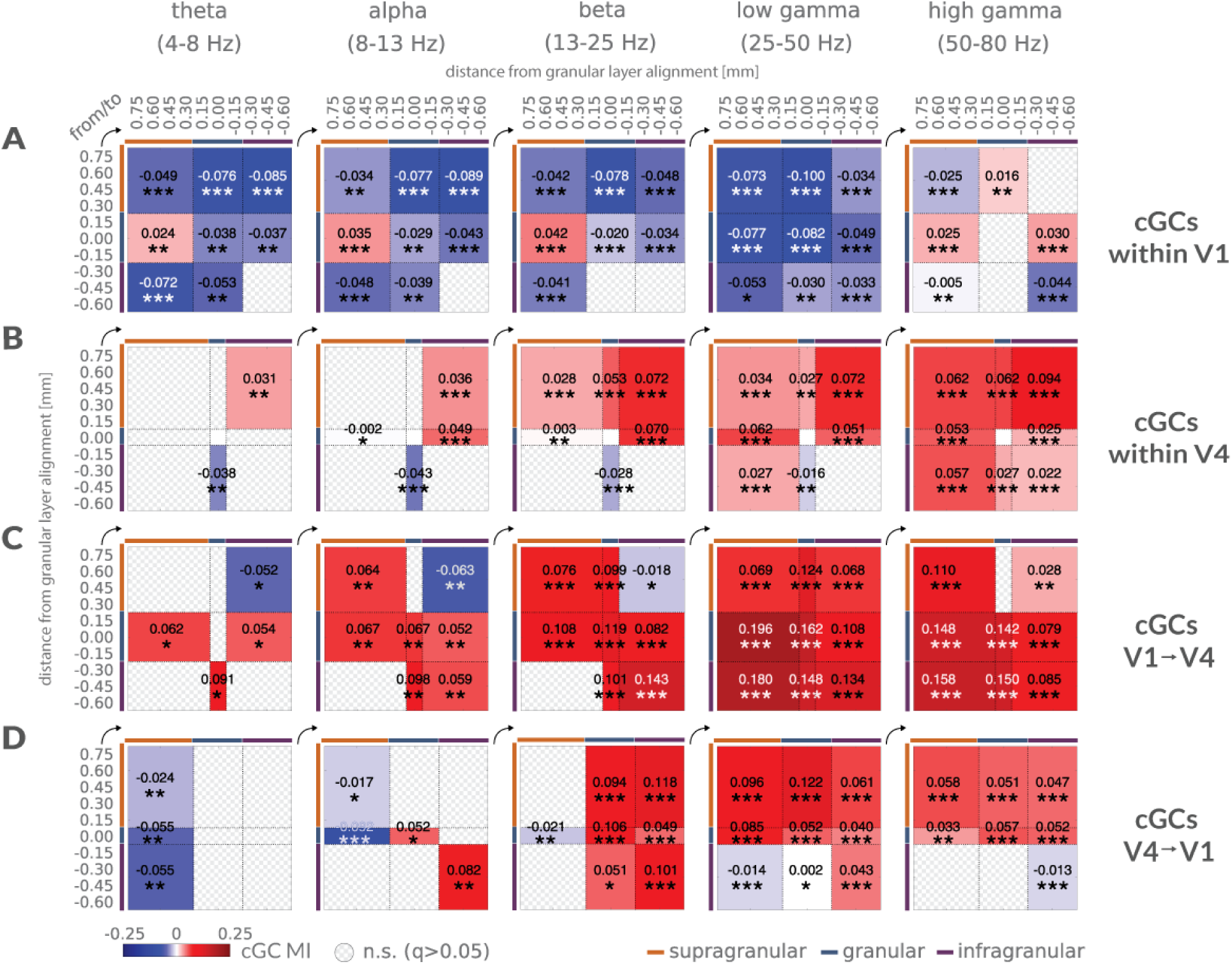
Attentional modulation of compartment-wise cGCs. **A)** Significant attentional modulation index of cGC (cGC MI) among compartment pairs within V1, (mean across sessions, pooled for the two monkeys), at different frequency bands (significance assessed via two-sided Wilcoxon signed rank tests, FDR corrected within frequency bands; * q≤0.05, ** q≤0.01, *** q≤0.001). **B)** Same as in A, but for V4. **C)** Same as in A, but for cGCs MIs from V1 to V4. **D)** Same as in A, but for cGCs MIs from V4 to V1.

Attentional modulation of cGCs in V4 was very different to the pattern seen in V1 (Figure 6B, 7B). Across all frequency ranges it increased from supragranular to infragranular layers but decreased from infragranular to granular layers for theta to low gamma frequencies. This could enable feedback information to flow prominently to lower areas (supragranular V4 to infragranular V4 and onwards to e.g. V2, V1), while at the same time limiting potentially inhibitory interactions (assuming infragranular layers communicate inhibitory prediction signals) on stimulus related processing (V4 infragranular to granular layers). In addition, downward communication (supragranular to infragranular) was increased by attention from theta to low gamma frequencies (Figures 6B, 7B). With an increasing oscillatory frequency, attentional modulation of cGCs between compartments increased, such that for high gamma frequencies, communication increases occurred across almost all compartments.

Despite the overall reduction of cGCs by attention within V1, its influence on V4 increased across frequency bands for most compartment comparisons (Figure 6C, 7C). In lower frequency bands, attention increased cGCs from V1 granular to all V4 layers (except for theta V1-V4 granular-granular interactions). However, in the theta to beta band V1 supragranular to V4 infragranular interactions were decreased. In the gamma frequency bands almost all V1 to V4 interactions were increased by attention.

cGCs from V4 to V1 were decreased by attention in the theta band from all V4 layers to V1 supragranular layers. In the alpha band significant decreases occurred from granular and supragranular V4 to supragranular V1, (Figure 6D, 7D). In the beta and low gamma band, attention increased V4 to V1 cGCs in a downwards direction (V4 supragranular to V1 granular and infragranular layer; from V4 granular to V1 granular and infragranular layer, and from V4 infragranular to V1 infragranular layer, Figures 6D, 7D). In the high gamma range attention increased cGCs from V4 supragranular and from V4 granular layers to all V1 layers, but decreased cGCs from V4 infragranular to V1 supragranular and infragranular layers.

These patterns of attentional modulations are summarized in a frequency dependent ‘influencer’ diagram in Figure 8. It shows the attention dependent reduction in cGCs across cortical layers and frequencies within V1, which nevertheless resulted in an increase in cGCs from area V1 to area V4. Feedback interactions were reduced by attention in the theta band, but mostly increased in the beta and gamma band. Within V4 cGCs were mostly increased in the beta and gamma bands. Some of these interactions are predicted by established theories of the frequency specificity interactions of feedforward and feedback connections, but many were in violation of established theory.

**Figure 8:**
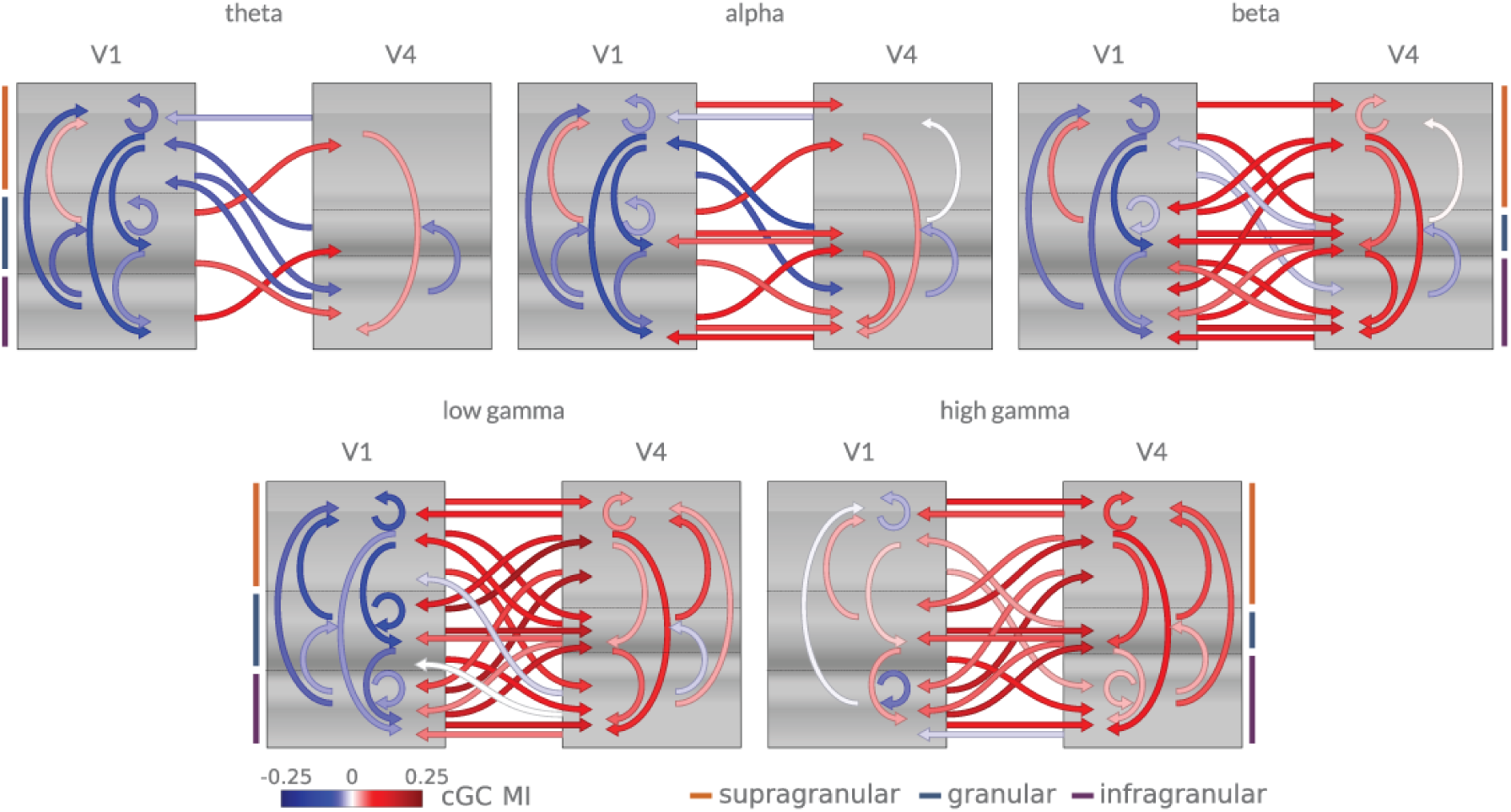
Main effects of attention on directed communication in different frequency bands. Arrows show significant attentional cGC modulation indices (cGC MI) (mean across sessions, pooled for the two monkeys), for the three laminar compartments (supragranular, granular, infragranular). Color indicates whether attention increases directed communication (red), or whether it decreases directed communication (blue), within and between areas.

## Discussion

Granger causal communication and its modulation by attention within and between areas V1 and V4 partly confirms, but also challenges current models of cortical processing. Within V1 dominant communication streams are directed towards supragranular cortico-cortical feedforward outputs. Conversely, in V4, dominant communication was bi-directional, with one stream of supragranular cortico-cortical feedforward flow, and a separate stream of supra- to infragranular feedback flow. Stimulus driven feedforward communication from V1 to V4 dominated in theta and gamma frequency ranges, with little layer specificity. Stimulus driven feedback communication from V4 to V1 dominated in the low frequency range. Surprisingly, attention to the receptive field generally reduced communication between cortical layers in area V1, with a notable exception for granular to supragranular communication. Within area V4, attention predominantly increased communication in beta and gamma frequency ranges. Despite the attention induced decrease of intra-areal V1 communication, attention increased feedforward communication from V1 to V4 across frequency bands. Attentional effects on feedback communication (V4 to V1) differed between frequency ranges. Theta and alpha communication decreased, while beta and gamma communication increased. Thus, feedforward interactions within and between cortical areas are neither limited to, nor dominant, in the gamma frequency range. Moreover, attention does not selectively increase gamma feedforward communication. Finally, feedback interactions between cortical areas, while dominant in the lower frequency range, are generally decreased by attention at low frequencies, but increased by attention in the gamma band.

For V1 LFPs, spectral power peak locations in the gamma range differed between attend RF and attend away conditions. The peak location for attend RF conditions resided at higher frequencies than for attend away conditions (∼3-4 Hz). An equivalent result using ECoG surface recordings has been interpreted as a shift towards higher gamma peak frequencies induced by attention (Bosman *et al.*, 2012). However, our comparison with steady state post stimulus gamma peak locations shows that attention keeps the peak gamma frequency closer to the stimulus induced gamma frequency, i.e. attention stops it from dropping. This difference in interpretation is important, as it speaks to the role of attention, and potential mechanisms involved. Attention affects normalization circuits, causing a concomitant increase in excitatory and inhibitory drive of the attended object/location (Carandini and Heeger, 2013; Lee and Maunsell, 2009; Ray *et al.*, 2013; Reynolds and Heeger, 2009; Sanayei *et al.*, 2015). This increases the power and the frequency of gamma oscillations (Gieselmann and Thiele, 2008; Ray and Maunsell, 2010; Ray *et al.*, 2013). Attention thus ensures that stimulus representations remain sensory input driven, and sustained responses remain elevated (Pooresmaeili *et al.*, 2010; Reynolds *et al.*, 1999; Reynolds and Heeger, 2009; Treue and Maunsell, 1999; Williford and Maunsell, 2006). Within a predictive coding framework (PC) (Bastos *et al.*, 2012; Kanai *et al.*, 2015; Rao and Ballard, 1999), this could be interpreted in two ways. First, if attention reduced prediction generation for attended locations/features/objects, then prediction error coding populations would respond more strongly to sensory stimuli, as these are less predicted. This in turn increases feedforward communication, which has been associated with gamma frequency oscillations (Bosman *et al.*, 2012; van Kerkoerle *et al.*, 2014; Von Stein *et al.*, 2000). Second, according to an extension of predictive coding that allows attentional signatures to arise naturally within the model (Feldman and Friston, 2010; Spratling, 2008), attention increases the precision of predictions, making neurons respond more strongly to hidden causes (sensory input). Gamma oscillations, as a signal of prediction errors (Bastos *et al.*, 2012; Bastos *et al.*, 2015) in superficial layers would thus be increased. Which of the two interpretations is correct remains to be determined.

We did not find consistent increases in gamma power with attention in V1 (only consistent differences in peak location were found). However, V4 gamma power, and peak location were increased in both monkeys, in line with previous reports (Bosman *et al.*, 2012; Buffalo *et al.*, 2011; Fries *et al.*, 2001; Gregoriou *et al.*, 2009; Vinck *et al.*, 2013). Prominent gamma oscillations and its modulation by attention have been argued to be largely confined to supragranular layers (Buffalo *et al.*, 2011; Maier *et al.*, 2010; Xing *et al.*, 2012). We did not find major differences in absolute gamma power, gamma power peak location, or attentional modulation of gamma power across supra-, granular, or infragranular layers in either V1 or V4. Using local bipolar referencing for all our analyses ensured that this was not an artefact of volume conduction. It shows that gamma frequency oscillations are not restricted to superficial layers, and they are thus unlikely a unique signature of feedforward interactions. Attention reduced oscillatory activity in theta and alpha bands in area V1 and V4, consistent with previous work (Bastos *et al.*, 2015; Bollimunta *et al.*, 2011; Brunet *et al.*, 2015; Buffalo *et al.*, 2011; Chalk *et al.*, 2010; Spyropoulos *et al.*, 2018; van Kerkoerle *et al.*, 2014). However, just as for gamma oscillations, these changes were not restricted to infragranular layers, but occurred across laminar compartments. These results equally question a strict separation between layer specific oscillatory frequency bands (Maier *et al.*, 2010; Spaak *et al.*, 2012; Xing *et al.*, 2012), and their potential association with feedforward and feedback signaling. They are more in line with recent reports about alpha sources across different modalities in primary sensory cortex (Haegens *et al.*, 2015).

### Communication across layers within and between areas

Interareal cGCs support the proposal that gamma and theta frequency interactions dominate in the feedforward direction (V1 to V4. Bastos *et al.*, 2015; Bosman *et al.*, 2012; Spyropoulos *et al.*, 2018; van Kerkoerle *et al.*, 2014), while alpha and beta frequency interactions dominate in the feedback direction (V4 to V1. Bastos *et al.*, 2015; Bosman *et al.*, 2012; Spyropoulos *et al.*, 2018; van Kerkoerle *et al.*, 2014). However, cGCs within areas deviated from this scheme in important aspects. While local feedback interactions from infragranular to granular layers and to supragranular layers were most prominent at low frequencies (Spaak *et al.*, 2012; van Kerkoerle *et al.*, 2014), strong and dominant low frequency cGCs from granular to supragranular layers occurred. Moreover, dominant gamma cGC intra-areal feedback direction occurred (from infragranular to granular and to supragranular) were present, which have been argued to label feedforward circuits (van Kerkoerle *et al.*, 2014). Thus, all cGCs in V1 dominate in a direction that targets the cortico-cortical output (supragranular) layers. This was the case for all frequencies, irrespective of the assumed role of oscillations in different frequency bands (Babapoor-Farrokhran *et al.*, 2017; Bastos *et al.*, 2015; Bonnefond and Jensen, 2013; Buschman *et al.*, 2012; Fries, 2015; Gregoriou *et al.*, 2009; Spaak *et al.*, 2012; Womelsdorf *et al.*, 2010). It suggests that V1 plays a key role as a distributor of feedforward information, with relatively less responsibility of feedback processing (as a consequence, it may have little effect in the generation of predictions. Bastos *et al.*, 2012; Feldman and Friston, 2010). The pattern changes slightly in V4, but it equally violates some key predictions about feedforward and feedback interactions. Namely, low frequency cGCs dominated in the feedforward direction (supra- to infragranular layers), while they dominated in the feedback direction in the gamma band (infra- to supragranular layers).

Attention to the RF reduced almost all cGCs within area V1, except for low frequency interactions from granular to supragranular layers. In the low gamma frequency band even those interactions were reduced, while most interactions were increased in the high gamma frequency band. The increase of cGCs from granular to supragranular layers is likely to boost feedforward output to other cortical areas, an expected effect given the increased efficacy demonstrated for feedforward spiking interactions and thalamocortical interactions with attention (Briggs *et al.*, 2013; Hembrook-Short *et al.*, 2019). If most intra-columnar feedback interactions served to compute context, while spatial attention boosts elementary processing (at the expense of contextual processing), then these cGC reductions are expected. Low frequency bands may predominantly play inhibitory roles (Bonnefond and Jensen, 2013; Haegens *et al.*, 2011; Spaak *et al.*, 2012). If these were reduced by attention, the increased firing rate seen in V1 in our and other studies (Hembrook-Short *et al.*, 2019; Herrero *et al.*, 2013; McAdams and Maunsell, 1999; Roelfsema *et al.*, 1998; Sanayei *et al.*, 2015; Wannig *et al.*, 2011) would be a natural consequence. Within the PC framework, it could be postulated that attention reduces the relative weight of predictions (although this is contrary to the proposal put forth by Feldman and Friston, 2010). Intuitively, attending to stimuli from the external world could mean re-shifting the balance from inferential to actuality processing, i.e. reducing the weight of internal priors. This would be achieved through reduction of feedback (local and interareal) and increase of feedforward processing. Such a re-shifting has been shown to be mediated by acetylcholine (Hasselmo and Bower, 1992; Roberts *et al.*, 2005), which plays an important role in attention (Dasilva *et al.*, 2019; Deco and Thiele, 2011; Herrero *et al.*, 2008; Roberts *et al.*, 2005).

The attentional modulation of cGCs in V4 differed radically from that in V1. Attention increased theta to beta band cGCs from supra- to infragranular layers and reduced theta to beta band cGCs from infra- to granular layers. In gamma bands almost all cGCs were increased. V4 is a major recipient of feedback from attentional signals originating in FEF (Gregoriou *et al.*, 2012; Gregoriou *et al.*, 2009; Moore and Armstrong, 2003; Moore *et al.*, 2003). The feedback is excitatory and predominantly targets excitatory cells in layer 2/3 (Anderson *et al.*, 2011). It could explain why cGCs originating from V4 supragranular layers show the most pronounced increases with attention. However, it does not explain why it occurs across all frequencies, if low frequency interactions label inhibitory interactions. Our data suggest that this association with inhibitory roles is debatable for the case of FEF-V4 interactions, as we do not expect attention mediated feedback to increase inhibition, The strong increases of cGCs between all layer compartments across frequency bands in V4 suggest that feedback and feedforward intracolumnar interactions within V4 do not strongly differentiate between frequencies.

Interactions from V1 to V4 were mostly increased by attention across frequency bands. Attentional increase was most profound in the gamma band, in line with the notion that gamma oscillations mediate feedforward communication (Bastos *et al.*, 2015; Bosman *et al.*, 2012; van Kerkoerle *et al.*, 2014). However, low frequency interactions were also increased, which questions the generality of imputing feedforward communication exclusively to the gamma band.

A structure involved in coordinating large scale network interactions is the pulvinar, which regulates cortical synchrony in an attention dependent manner, particularly in the low frequency range (Saalmann *et al.*, 2012). However, pulvinar also affects oscillatory activity in the gamma frequency range in V4 (Zhou *et al.*, 2016). The changes seen for V1 to V4 cGCs in the low frequency range could be mediated through cortico-pulvinar-cortical interactions (Sherman *et al.*, 2002; Shipp, 2003). This might also explain the relative absence of layer specificity in cGC interactions between V1 and V4, irrespective of their direction.

Attention reduced communication from V4 to V1 in the theta band, and most strongly increased cGC in the beta band. However, strong increases also occurred in the gamma band, demonstrating that feedback interactions also operate strongly in the gamma band. V4 to V1 cGCs equally did not show strong laminar specificity. While this could be a consequence of subcortical routing (Sherman *et al.*, 2002; Shipp, 2003), it could also be a consequence of a termination pattern of V4 feedback that predominantly targets layer 1 dendritic spines through excitatory synapses (Anderson and Martin, 2006). These terminations can thereby influence pyramidal cells across supra- and infragranular layers. The predominance of excitatory connections on pyramidal cell dendrites is not consistent with the proposal that predictions generated at higher cortical levels act through di-synaptic inhibition for messages passing to lower areas (Bastos *et al.*, 2012).

A recent theory of ‘predictive routing’ (Bastos *et al.*, 2020) proposed that low frequency feedback prepares feedforward pathways, by inhibiting gamma and spiking activity associated with predicted inputs. A reduction in feedback (prediction) signals would thus cause disinhibition. Our results align, but also argue for an extension of this predictive routing scheme. We argue for different hierarchies of prediction generation, some are automatic (e.g. surround suppression, basic contour integration, contrast normalization), while others are associated with higher cognitive functions (e.g. working memory, feature search, spatial attention, value estimation). We also propose that these to some extent employ different feedback networks. Automatic prediction generation mostly works within connections that affect non-classical receptive field interactions. This would explain why cooling of higher level areas results in reduced surround suppression (Hupé *et al.*, 1998), i.e. upon cooling, higher areas cannot pass predictions to lower areas. Inhibition is thus reduced and prediction error (or to use different words, sensory coding) signaling will be large. On the other hand, interactions between neurons sharing classical receptive field (cRF) locations counterbalance the prediction coding, i.e. they are predominantly excitatory. This explains why cooling of higher cortical areas results in reduced cRF responses (Hupé *et al.*, 1998). It is these cRF routes that might be exploited by higher cognitive functions, which through a separate form of feedback generate biased competition, and simultaneously serve to suppress automatic Bayesian inference (PC). Our data of attention induced increased feedforward, but decreased feedback communication within V1, increased feedforward and feedback cGCs within V4, and increased bidirectionally communication between V1 and V4 (with overlapping cRFs) across most frequency ranges support such a proposal.

## Acknowledgments

Funded by Wellcome Trust 093104 (JvK, MB, AT), MRC MR/P013031/1 (JvK, AT), NIH Brain Initiative R01 NS108410 and U19 NS107464U19 (SP) and Simons Foundation SFARI Explorer grant 602849 (SP).

## Author contributions

Demetrio Ferro: Data analysis and analysis methods, data curation, manuscript writing, visualization

Michael Boyd: data acquisition,

Jochem van Kempen: data acquisition, data analysis, manuscript review,

Stefano Panzeri: Data analysis and analysis methods, manuscript writing and review, supervision, funding acquisition,

Alexander Thiele: Conceptualization, data acquisition, resources, manuscript writing and review, supervision, funding acquisition.

## Methods

### EXPERIMENTAL PROCEDURES

#### Animals and procedures

We simultaneously recorded from visual areas V1 and V4 of two adult male rhesus macaque monkeys (Macaca mulatta, 10-11 years of age), while they performed a sustained top-down, feature-guided, visuospatial attention task. Experimental procedures were in line with the Directive 2010/63/EU of the European Parliament and of the Council of the European Union, the Guidelines for Care and Use of Animals for Experimental Procedures from the National Institute of Health, the Policies on the Use of Animals and Humans in Neuroscience Research from the Society for Neuroscience, and the UK Animals Scientific Procedures Act. Animals were motivated to engage in behavioural tasks through fluid control at levels that do not affect animal physiology and have minimal impact on psychological wellbeing (Gray *et al.*, 2016).

#### Surgical preparation

Animals were implanted with a head post and recording chambers over area V1 and V4 under sterile conditions and general anaesthesia. Surgical procedures and postoperative care conditions have been described in detail previously (Thiele *et al.*, 2006).

#### Behavioral paradigm

Monkeys were trained to comfortably sit in a primate chair while being head stabilized by the cranial head holder. Stimuli were presented on a cathode ray monitor (22’’ CRT, 120Hz, 1280×1024 pixel resolution) placed at 54 cm distance to the monkey’s eyes. Eye position was calibrated and monitored by an eye tracking system operating at a sampling rate of 220Hz. Stimulus presentation and behavioral control was handled by Remote Cortex 5.95 (Laboratory of Neuropsychology, National Institute for Mental Health, Bethesda, MD).

#### Attention Behavioral Task

Monkeys had to touch a lever for the appearance of a centrally placed fixation spot. Thereafter they had to direct their gaze at a fixation point (FP) positioned at the center of the CRT screen, with a fixation window of ± 0.7°-1.5° of visual angle (DVA) throughout the trial duration.

500 ms after fixation onset monkeys were presented with three colored, moving grating stimuli positioned equidistant from the FP. One stimulus was centered on the receptive field (RF) of recorded cells in V1, the other two were positioned outside (at locations OUT_1_ and OUT_2_). The RFs of recorded cells were mapped at the beginning of each experimental session (see below).

630 - 960 ms after stimulus onset (random delay, uniformly distributed, 1 ms steps) a colored cue was presented at FP. The color of the cue instructed the monkey to monitor the stimulus of matching color (e.g. a red cue instructed the animal to monitor the red visual stimulus) for a change in luminance contrast and ignore changes at the other stimulus locations.

After a random delay, the three stimuli started to sequentially dim in a pseudo-random order. Delays for subsequent dimmings ranged between 1160 – 1820 ms (the first dimming could occur 1160 - 1820 ms after cue onset, the second dimming could occur 790 - 1120 ms after the first dimming, etc.).

During the entire trial period monkeys had to keep fixating the FP. Upon cued stimulus dimming, monkeys had to release the touch bar within 600 ms to receive a fluid reward. Figure 1A graphically shows the time course of a sample trial of the main behavioral task. The grating stimuli had a diameter between 2 to 4 DVA, adjusted in accordance with the size and eccentricity of the recorded RFs. Their spatial frequency was 1.5 cycles/DVA, with a temporal frequency of 1 cycle/s (in sessions where they moved) and an orientation of 30°. The stimulus color at a given location was fixed (red, green or blue) for trials of the same session but randomized across sessions to cover all the 6 possible color configurations. In the same way, the cue color (red, green or blue), the order of dimming of the three stimuli (6 possible dimming orders), as well as the direction of movement of the grating stimuli (2 possible opposite directions, where applicable) were pseudo-randomized across trials to cover all possible task configurations.

Thus, there were 36 conditions total, which comprised a so-called cycle. In each cycle all 36 conditions would occur at least once, selected on a random basis. If the monkey performed the trial correctly, the condition was removed from the cycle pool. If the trial was not completed correctly, the condition was reinserted into the cycle pool, and would be reselected on a random basis, until all conditions had had been performed correctly.

#### Electrophysiological recordings

Electrophysiological recordings were performed using passive laminar probes with 16 recording contacts, inter-contact spacing of 150 μm (ATLAS Neuroengineering, Belgium). The laminar probes were inserted perpendicularly to the cortical surface with the support of a hydraulic micromanipulator (NARISHIGE MO-97A, Japan). All contacts were initially referenced to a wire positioned either in the V1 chamber, or in the V4 chamber.

Data from the two chambers were simultaneously recorded using a digital acquisition and control system (Digital Lynx, Neuralynx, USA) with a sampling frequency of 32556 Hz (∼32 kHz), at 24 bits.

The data were collected over 62 sessions (34 for monkey 1, 28 for monkey 2), yielding a total of 35744 correct trials (15892 in monkey 1, 19852 in monkey 2). These were out of 36912 total trials (16698 in monkey 1, 20214 in monkey 2), where monkeys kept fixation, yielding a behavioral performance of 95.17% correct for monkey 1, and 98.21% correct for monkey 2.

#### Receptive Field Mapping

Prior to starting the attention paradigm, the location and size of the RF was measured by a reverse correlation method (Gieselmann and Thiele, 2008).

From this, RF maps were initially estimated online to determine the stimulus locations in the attention paradigm. Offline RF analysis was done based on spike sorted single units, on thresholded multi-unit activity, as well as based on local population activity (envelope multi-unit activity, MUA_E_, Supèr and Roelfsema, 2005) using a time window from 50-130 ms after RF mapping stimulus onset.

### OFFLINE DATA ANALYSIS

#### Electrophysiological data analysis

Signals were extracted in time windows relative to task-related events: after stimulus onset (0 to 503.25 ms, 512 data points see below for LFP sampling frequency), after cue onset (0 to 503.25 ms, 512 data points see below for LFP sampling frequency) and before dimming times (503.25 ms before each of the three subsequent dimmings, 512 data points see below for LFP sampling frequency). Baseline activity time window started 200 ms before stimulus onset and covered up to 30 ms after stimuli onset.

Data were replayed offline, sampled with 16-bit, band-pass filtered at 0.5-300 Hz and down sampled by a factor of 32 to a sampling frequency F_s_ =1017.375 Hz to obtain local field potential (LFP) data. Spiking Activity was accessed by band-pass filtering between 600 and 9000 Hz, then further analyzed both at the level of multi-unit activity by extracting the Multi-Unit Activity Envelope (MUA_E_), and by sorting single-unit spiking waveforms for RF mapping. Spikes were sorted manually using SpikeSort3D (Neuralynx).

#### Multi-Unit Activity Envelope and Signal to Noise Ratio

MUA_E_ was computed as described by (Supèr and Roelfsema, 2005). The higher frequency signal component (600-9000 Hz) was down sampled by a factor of 4, full-wave rectified, low-pass filtered at 500 Hz (Butterworth, zero-phase digital filter of order 5), and further down sampled by a factor of 8 to a frequency of 1017.375 Hz.

Signal to Noise Ratio (SNR) computation was performed on MUA_E_ signals in n = 8sliding time windows of length 50 ms shifted every 10 ms from 30 to 150 ms after stimuli onset. The SNR is computed as the maximum average magnitude of baseline corrected MUA_E_ signal across time windows, i.e. SNR = max_n_{(⟨s_n_(t)⟩ − ⟨b(t)⟩)/σ_b_}, where ⟨s_n_(t)⟩ is MUA_E_ average within time window n, and ⟨b(t)⟩ and σ_b_ are respectively the baseline mean and standard deviation.

#### Laminar alignment

Laminar signals from different experimental sessions were aligned to layer IV of both V1 and V4. Layer IV was identified for each session as the earliest current sink across laminae using current source density (CSD) of LFPs, and by analyzing the shortest latency of the stimulus evoked MUA_E_ response. Based on their distance from reference coordinate, signals from the corresponding recording channels were assigned to three main laminar compartments: supragranular, granular and infragranular. For V1, channels above the reference channel at distances of 0.25 - 1 mm were labelled as supragranular, channels above or below the reference channel within 0.25 mm were labelled as granular, and channels below reference at distance range 0.25 - 0.75 mm were labelled as infragranular. For V4, channels above the reference channel in the range 0.1 - 1 mm were labelled as supragranular, channels within 0.1 mm above or below the reference channel were labelled as granular, channels below the reference channel at 0.1 - 0.75 mm were labelled as infragranular.

#### Current Source Density analysis

The current source density (CSD) signal was obtained by applying the spline inverse CSD (iCSD) method (Pettersen *et al.*, 2006). Starting from the direct equation for the field potential **Φ** generated by a point source **C** positioned at the origin of an isotropic medium **Φ = F ⋅ C**, the iCSD was estimated by inversion of the conduction matrix **F** as **Ĉ = F^−1^ ⋅ Φ**. The coefficients of F were computed by electrostatic field equations for point sources by assuming that they are evenly distributed within isotropic cylindrical discs of finite radius R, and by assuming smooth CSD variation along depth dimension. CSD variation along depths was approximated by cubic splines interpolation. In our computations we assumed a disc radius R =500 μm (Mountcastle, 1957), and we used conductance σ = 0.4 S/m (Logothetis et al., 2007). The conductance term could affect the magnitude of iCSDs but not their spatial profile. The iCSD was filtered by a Gaussian filter with standard deviation 200 μm along the depth dimension.

#### Response Latency analysis

The method used to compute stimulus response latency followed the formulation by Roelfsema *et al.* (2007), i.e. by assuming that stimulus responses arises at random times with a Gaussian distribution, and that a fraction of the response function dissipates exponentially after reaching a peak magnitude. On our data, the response function was estimated by fitting the 150 ms baseline-corrected post-stimulus MUA_E_ signals to a distribution f(t) consisting in the sum of ex-Gaussian and cumulative Gaussian functions:

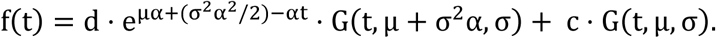

The parameters μ and σ respectively match the mean and standard deviation of the response function onset time when considering the response as non-dissipating. The parameter α is the dissipation rate, and the parameters c and d act as weighting factors for the response magnitude and dissipation terms. The functions G(t, μ^′^, σ^′^) are cumulative density functions of a generic normally distributed variable with mean μ′ and standard deviation σ′.

Response latency is computed as the smallest time delay allowing to achieve 33% of the peak in the response magnitude 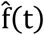 estimated by least-square error minimization. In symbols, we computed latency as: 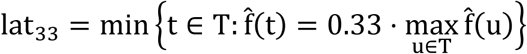. To reduce computational cost of the least-square fit procedure, the empirical MUA_E_ response was smoothed by a moving average filter with length 5 samples, covering ≈5 ms.

#### Trials and Channels inclusion criteria

All our analyses only included trials with behaviorally successful outcome. To correct for eventual artifacts, which could be due to transient drifts of the probe, possibly caused by slight movements of the animal, we set a signal thresholding rule for trials selection. Trials were discarded if the baseline normalized signal energy was higher than the energy of a signal with magnitude 20 times bigger than baseline, i.e. if the signal 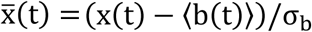 had energy 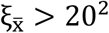, where 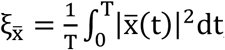, x(t) is the LFP/MUA_E_ signal in any of the task-relevant time windows, ⟨b(t)⟩ and σ_b_ are the signal mean and standard deviation at baseline.

Applying this thresholding rule led to the rejection of 2.1% of the trials, hence to the selection of 34992 out of 35744 behaviorally correct trials (15468 were from monkey 1, 19524 for monkey 2).

In all analyses we ensured to use equal amount of trials per attentional condition (RF, OUT_1_, OUT_2_) by random sub-selecting trials in each session so that the amount of trials per condition was equal to the minimum amount of trials available in the three conditions.

To prevent signal contamination due to common grounding or strong remote signal sources, the signals for each electrode contact from the two cortical areas were locally referenced via bipolar differentiation. The signal from depth z_i_ was replaced by the difference between signals at depths z_i+1_ and z_i−1_, as if it was recorded by a virtual electrode located at intermediate depth between its two neighboring contacts. This procedure could not allow us to consider the two most outer channels (as they could not be re-referenced with respect to their neighbor channels), but this was often not problematic as the channels located at outer positions were usually outside the grey matter of the targeted cortical areas. In addition, the quality of signals recorded from any of the channels was determined by the computation of SNR, and we only included channels with SNR ≥ 3. This resulted in data included from 481 channels for V1 (224 in monkey 1, 257 in monkey 2) and 531 channels for V4 (306 in monkey 1, 225 in monkey 2).

#### Spectral Power

The estimation of LFP signal power across frequencies was performed using a multi-tapering approach (Thomas, 1982). We used the Chronux toolbox developed by Mitra and Bokil (2008). We set the tapering to K = 3 Slepian waveforms with time-bandwidth product TW = 2 (T = N/F_s_ ≈ 500 ms, W ≈ 4 Hz). The LFP spectral power S_i_(λ) was normalized for each λ ∈ [0, F_s_/2] to baseline spectral power (minus trial-averaged baseline power, divided by the standard deviation of baseline power).

#### Spectral Coherence

The relationship between the spectral components of LFP signals recorded from multiple channels was quantified in terms of spectral coherence. This measure is computed by means of the cross-spectrum power density 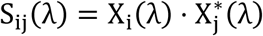, involving the spectral representations X_i_(λ) and X_j_(λ) of signals in channels i and j. The spectral coherence is defined as C_ij_(λ) = |S_ij_(λ)|^2^ /|S_i_(λ) ⋅ S_j_(λ)|, λ ∈ [0, F_s_/2]. The values assumed by C_ij_(λ) are in the range [0,1], where 0 means that the frequency components of the two signals are completely unrelated, and 1 means the two signals have perfectly linear relationship at given frequency component. The terms S_i_(λ), S_j_(λ), and S_ij_(λ) were computed with the use of the Chronux toolbox via multi-taper estimation (using K = 3 Slepian sequences, TW = 2).

#### Time-frequency spectral modulation

The spectral characterization was also performed in the time/frequency domain. LFP spectral power and coherence were both computed by using sliding time windows of duration 503.25 ms (N =512 time points), shifted in time every 20 ms to cover 1000 ms before the time of first stimulus dimming. The spectral resolution was Δf = F_s_/N ≈ 2Hz and temporal resolution was Δt =20 ms.

#### Granger Causality Analysis

We measured directed causal communication between LFPs recorded at different contacts by using Granger causality (GC). We analyzed GC in a 503.25 ms time window (512 time points at 1017.375 Hz sampling rate) preceding the first dimming time. To reduce computational time, the signals were down sampled to 128 time points at a sampling frequency of 254.34 Hz.

In its original formulation (Granger, 1969; Geweke, 1982), the GC between two times series Y(t) and X(t) is computed by fitting a multivariate vector autoregressive model (MVAR) with finite memory p. The fit consists in estimating the linear interaction coefficients **A**_k,k=1…p_ by least squares regression, yielding residual fit error of mean zero and covariance **Σ**.

Spectral GC is characterized at each frequency λ ∈ [0, F_s_/2] via the cross-spectral density matrix **S**(λ) and the MVAR transfer function matrix **H**(λ) yielding factorization **S**(λ) = **H**(λ) ⋅ **Σ** ⋅ **H**(λ)^*^ (Geweke, 1982). Spectral GC is then defined as:

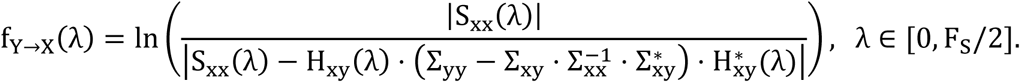

To provide a more refined measure of the communication between time series of LFPs in channels Y and X, in our analysis we computed the GC between Y and X conditioned on z (called here Conditional GC, with acronym cGC).

This more refined measure discounts the possible confounding effect of time-lagged interactions mediated by activity of other recorded channels **Z** = {Z_1_, …, Z_m_} rather than direct communication between the two considered nodes Y and X (see below for details of our infomax partial conditioning choice of the m channels **Z**).

Following the derivation by Geweke (1984), cGC f_Y→X|z_ was computed by first applying a reduced least-square autoregression to the time series X, **Z** only, yielding residual error time series X^†^, **Z**^†^. Then, cGC was defined via the identity: 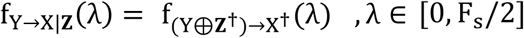, allowing to express cGC with the original definition as unconditional GC between the variables 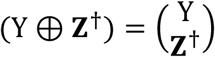 and X^†^.

In our analysis, cGC was computed by the ‘MVGC’ method based on the computation of MVAR autocovariance sequences via Yule-Walker equations using the ‘MVGC toolbox’ by Barnett and Seth (2014). The magnitude of spectral GCs did not qualitatively vary when using alternative methods such as matrix partitioning (Chen et al., 2006), nonparametric spectral factorization (Dhamala *et al.*, 2008), or time reversed GC (Vinck et al., 2015).

The stationarity of LFPs, an important check for the application of Granger analyses, was assessed by ensuring that the MVAR characteristic polynomial 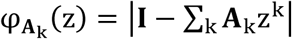 was invertible within unit disc, i.e. that 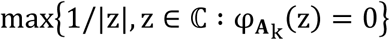 was always < 1 (Barnett and Seth, 2014).

An important aspect of the analysis is the choice of which and how many (*m* parameter) the channels **Z** are chosen for conditioning in cGC. Conditioning on all available channels complementary to Y and X (m = ALL) (‘full conditioning’) might suffer from lack of sufficient data to estimate all autoregressive models needed for this calculation. In addition, full conditioning would make cGC unevenly scaled (since the number of available channels could vary across sessions), and regressing out channels **Z** within the same laminar compartments of Y or X would likely discount genuine interactions, because of the stronger correlations between channels within the same compartment due to volume conduction or other effects. Following Marinazzo et al. (2012), we thus applied the infomax partial conditioning strategy. We considered for conditioning only channels outside the laminar compartments of channels X and Y.

We then chose **Z** to be the m channels with the highest mutual information with Y and X (cGCs for different m are shown in Supplementary Figure S5). Mutual information (Shannon, 1948) between channel pairs was computed on the Hilbert envelope of LFP time series demodulated in 2 Hz frequency bins, then integrated in time and frequency. We used the ‘Information toolbox’ (Magri *et al.*, 2009) for mutual information estimation, and the method by Panzeri and Treves (1996) for subtracting the limited sampling bias.

We chose the free parameters of the conditioning (the number and identity of conditioning channels) as the ones with the best Akaike Information Criterion (AIC) and the autoregression coefficient of determination R^2^ (Supplementary Figures S5A-D). The two indices were adjusted to the size of the data sample to AICc (Hurvich and Tsai, 1989) and 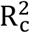 (Theil, 1961) in order to prevent from data overfitting, though the correction did not affect the results much. The optimization of AICc and 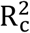 led us to the selection of an MVAR model with memory p=10 (≈40 ms), and conditioning variable z made by m = 2 channels outside the compartments of X and Y, achieving average R^2^=0.8, s.e.m.= 0.0006 (p=10 and m=2 (out) in Figure S5).

To estimate the statistical significance of the empirical cGCs and to exclude any possible residual limited sampling biases, we recomputed cGCs after randomly shuffling the data across trials (we used 100 different shuffles for each directed channel pair). The significance of empirical cGC magnitudes was then assessed by setting a significance threshold equal to the 95^th^ percentile of shuffled cGCs (Chen et al., 2006).

#### Attentional Modulation Index

To investigate the effects of attention, we compared results for the trials where attention was directed towards RF visual location against the ones where it was directed at outside locations OUT_1_, OUT_2_. Since the LFP spectral characterization for these two latter cases did not show prominent differences, we combined them in a single attend OUT condition by random subsampling an equal number of trials with condition OUT_1_ and OUT_2_. The modulation index (MI) for the measure F (spectral power or cGC) was defined as F_MI_ = (F_RF_ − F_OUT_)/(F_RF_ + F_OUT_).

#### Statistical tests and significance

In all our analyses, the significance of the difference in spectral power, coherence, or cGCs (e.g. between time windows [before stimuli onset and after stimuli onset], attentional conditions [attend RF vs attend OUT], directionality of cGCs (f_X→Y|z_ vs f_Y→X|z_)), as well as the significance of attentional modulation indices (F_MI_), were tested across experimental sessions by two-sided Wilcoxon signed rank tests (Wilcoxon, 1945). The p-values were corrected for False Discovery Rate (FDR) at q = 0.05 (Benjamini and Hochberg, 1995).

