## Supplementary Materials for "Directed information exchange between cortical layers in macaque V1 and V4 and its modulation by selective attention"

### SUPPLEMENTAL INFORMATION

#### Data acquisition hardware

- Electrode probe (16 recording contacts, inter-contact spacing of 150 µm, model E16R-150-S1-L10, brand Atlas Neuroengineering, Belgium);
- Hydraulic manipulator (model MO-97A, brand NASHIRIGE, Japan);
- Preamplifier (model HS36, Neuralynx, USA);
- CRT Monitor (size 22”, resolution 1280x1024 pixels, frame rate 120 Hz, model HM204DTA, brand Iiyama, Japan);
- ViewPoint Eye tracker (model BHU03, brand Arrington research, USA);
- Digital acquisition tool (Digital Lynx: 32 channels, sampling frequency of 32556 Hz digital quantization resolution of 24 bits).

#### Data acquisistion software

- Stimuli presentation and behavioral control: Remote Cortex 5.95 (Laboratory of Neuropsychology, National Institute for Mental Health, Bethesda, MD).
- Raw signals extraction/filtering: Cheetah 5.6.3 (Neuralynx Inc., USA).

#### Data Analysis software

- MATLAB® 2018b (Mathworks Inc.,USA);
- Neuralynx MATLAB-Netcom Utilities 6.0.0 (Neuralynx Inc., USA);
- Chronux Toolbox 2.12 chronux.org; (Mitra and Bokil, 2008);
- MVGC Toolbox (Barnett and Seth, 2014).
- Mutual Information toolbox (Magri *et al.*, 2009)

#### Color specifications

| Color | color code [R, G, B] | | | Luminance [cd/m$^2$] |
|---|---|---|---|---|
| **Gray (background)** | 45 | 45 | 45 | 0.8 |
| **Red** | 220 | 0 | 0 | 12.8 |
| **Green** | 0 | 135 | 0 | 12.9 |
| **Blue** | 60 | 60 | 255 | 12.2 |
| **Dimmed red** | 140 | 0 | 0 | 4.2 |
| **Dimmed green** | 0 | 90 | 0 | 4.6 |
| **Dimmed blue** | 30 | 30 | 180 | 4.6 |


Barnett, L., Seth, A.K. (2014) The MVGC multivariate Granger causality toolbox: A new approach to Granger-causal inference. *Journal of Neuroscience Methods*. 223, 50–68.

Magri, C., Whittingstall, K., Singh, V., Logothetis, N.K., Panzeri, S. (2009) A toolbox for the fast information analysis of multiple-site LFP, EEG and spike train recordings. *BMC Neuroscience*. 10.

Mitra, P.P., Bokil, H. (2008) *Observed Brain Dynamics*. Oxford University Press.

**Sessions, trials and behavioral performances.** Number of sessions for data recorded in V1 and in V4 and number of sessions where data were recorded simultaneously in both V1 and V4. Total number of task trials performed, number of task trials where attended stimulus was correctly reported, percentage of total trials resulting in correct behavior.

| | monkey 1 | | | monkey 2 | | | monkeys 1:2 |
|---|---|---|---|---|---|---|---|
| | **V1** | **V4** | **V1&V4** | **V1** | **V4** | **V1&V4** | **V1&V4** |
| Sessions | 34 | 35 | 34 | 30 | 30 | 28 | 62 |
| Correct trials | 15892 | 16301 | 15892 | 21632 | 21383 | 19852 | 35744 |
| Error trials | 806 | 818 | 806 | 443 | 384 | 362 | 1168 |
| Total trials | 16698 | 17119 | 16698 | 22075 | 21767 | 20214 | 36912 |
| Performance (%) | 95.17% | 95.22% | 95.17% | 97.99% | 98.24% | 98.21% | 96.84% |
| Missed (bar not released) | 588 | 596 | 588 | 526 | 508 | 526 | 1114 |
| Missed (fixation break) | 1407 | 1460 | 1407 | 3397 | 3362 | 3397 | 4804 |

**Selection of trials based on LFP signal energy.** Number of trials selected, and percentage of trials rejected in V1 and V4 by applying artefact removal criteria based on LFP signals energy. Number of simultaneous trials selected for V1 and V4 once the trials selection was applied to V1 and V4.

| | monkey 1 | | | monkey 2 | | | monkeys 1:2 |
|---|---|---|---|---|---|---|---|
| | **V1** | **V4** | **V1&V4** | **V1** | **V4** | **V1&V4** | **V1&V4** |
| Number of selected trials | 15594 | 16011 | 15468 | 21312 | 21120 | 19524 | 34992 |
| Percentage Rejected | 1.88% | 1.78% | 2.67% | 1.48% | 1.23% | 1.65% | 2.1% |

**Selection of channels based on SNR threshold.** Total number of channels and average number of channels for each session in V1 and V4 selected by SNR ≥ 3 thresholding rule. Total number of channels selected, and average number of channels selected for each session in each V1 and V4 laminar compartment following laminar depth assignment.

| | | monkey 1 | | monkey 2 | |
|---|---|---|---|---|---|
| | | **V1** | **V4** | **V1** | **V4** |
| SNR ≥3 | Total channels (chs/sessions) | 224 (6.59) | 306 (8.74) | 257 (8.57) | 225 (7.50) |
| | **Supragranular chs (chs/sessions)** | **53 (1.56)** | **158 (4.51)** | **89 (2.97)** | **109 (3.63)** |
| | **Granular chs (chs/sessions)** | **107 (3.15)** | **42 (1.20)** | **98 (3.27)** | **29 (0.97)** |
| | **Infragranular chs (chs/sessions)** | **64 (1.88)** | **106 (3.03)** | **70 (2.33)** | **87 (2.90)** |

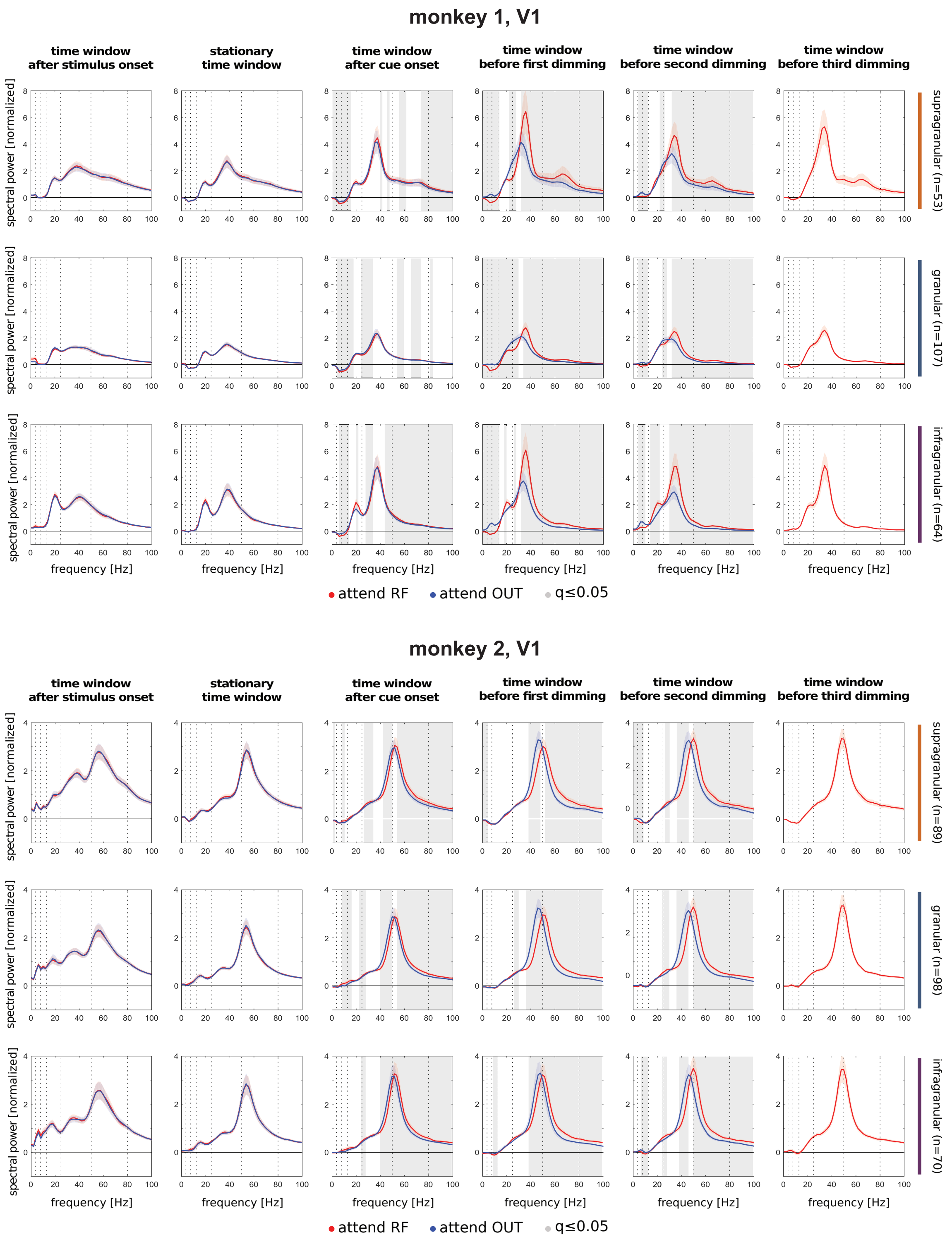


**Figure S1: Spectral power of V1 LFPs during the attentional task execution.** Panels in each column show baseline-normalized spectral power (mean +/- S.E.M across depths, sessions) computed in 503.25 ms (512 time points) task-related time windows (post-stimulus time window: 0 to 503.25 ms after stimulus onset; stationary time window: 200 to 703.25 ms after stimulus onset; 0 to 503.25 ms after cue onset; time windows before dimmings: -503.25 to 0 ms respectively before first, second and third dimming), in the two attentional conditions (attend RF, attend OUT). Baseline time window is set to -203.5 to 0 ms before stimulus onset. Panels with spectral power before second and third dimming just include trials with unchanged contrast in RF, implying that stimulus in RF location did not dim at previous times. Panels in each row refer to different laminar compartiments (supragranular, granular, infragranular), for monkey 1 (three top rows) and monkey 2 (three bottom rows). Dashed lines report the spectral bands selected (theta 4-8 Hz, alpha 8-13 Hz, beta 13-25 Hz, low gamma 25-50 Hz, high gamma 50-80 Hz), gray shaded background is for significant differences between attentional conditions (two-sided Wilcoxon signed rank tests, FDR corrected, q≤0.05).

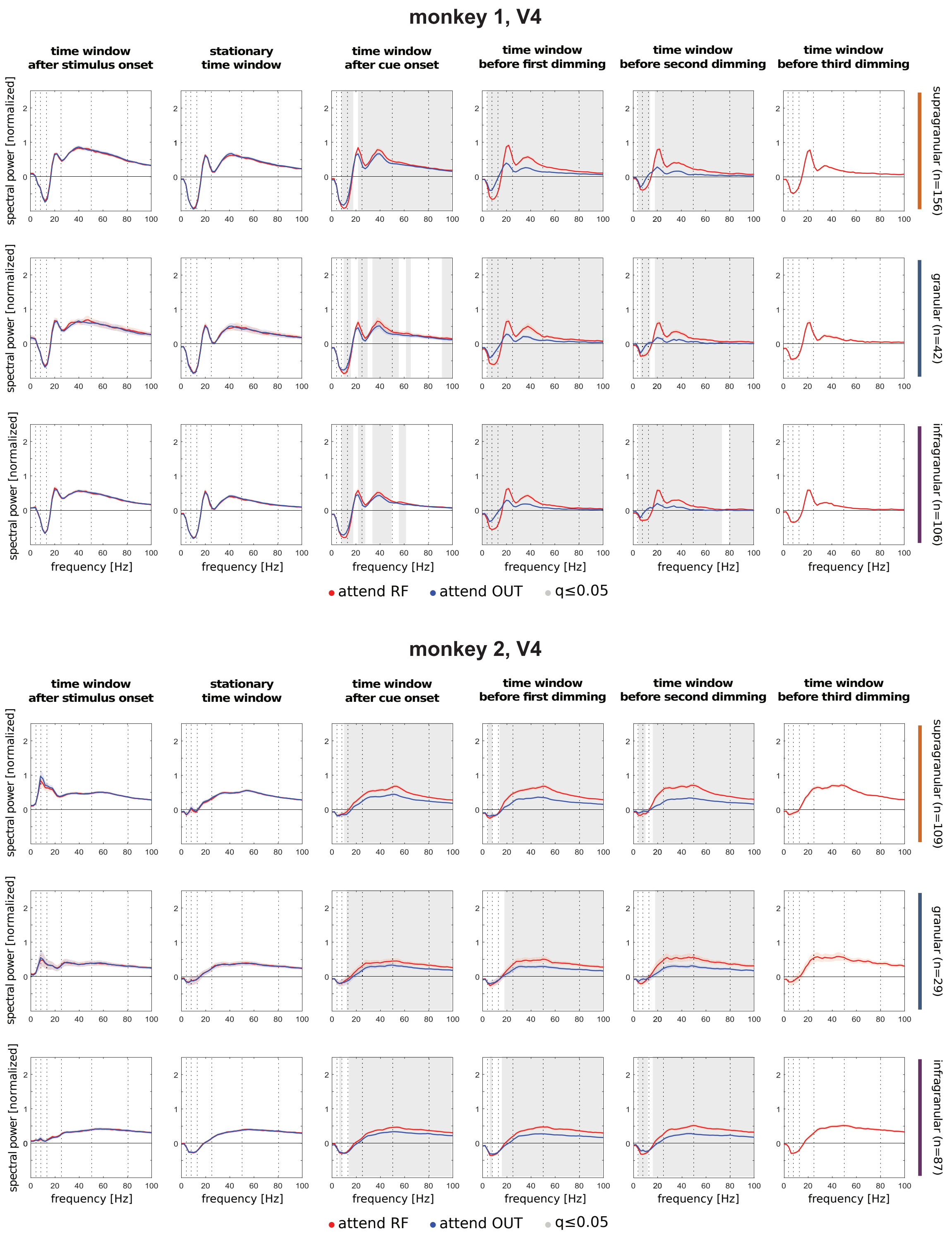


**Figure S2: Spectral power of V4 LFPs during the attentional task execution.** Panels in each column show baseline-normalized spectral power (mean +/- S.E.M across depths, sessions) computed in 503.25 ms (512 time points) task-related time windows (post-stimulus time window: 0 to 503.25 ms after stimulus onset; stationary time window: 200 to 703.25 ms after stimulus onset; 0 to 503.25 ms after cue onset; time windows before dimmings: -503.25 to 0 ms respectively before first, second and third dimming), in the two attentional conditions (attend RF, attend OUT). Baseline time window is set to -203.5 to 0 ms before stimulus onset. Panels with spectral power before second and third dimming just include trials with unchanged contrast in RF, implying that stimulus in RF location did not dim at previous times. Panels in each row refer to different laminar compartiments (supragranular, granular, infragranular), for monkey 1 (three top rows) and monkey 2 (three bottom rows). Dashed lines report the spectral bands selected (theta 4-8 Hz, alpha 8-13 Hz, beta 13-25 Hz, low gamma 25-50 Hz, high gamma 50-80 Hz), gray shaded background is for significant differences between attentional conditions (two-sided Wilcoxon signed rank tests, FDR corrected, q≤0.05).

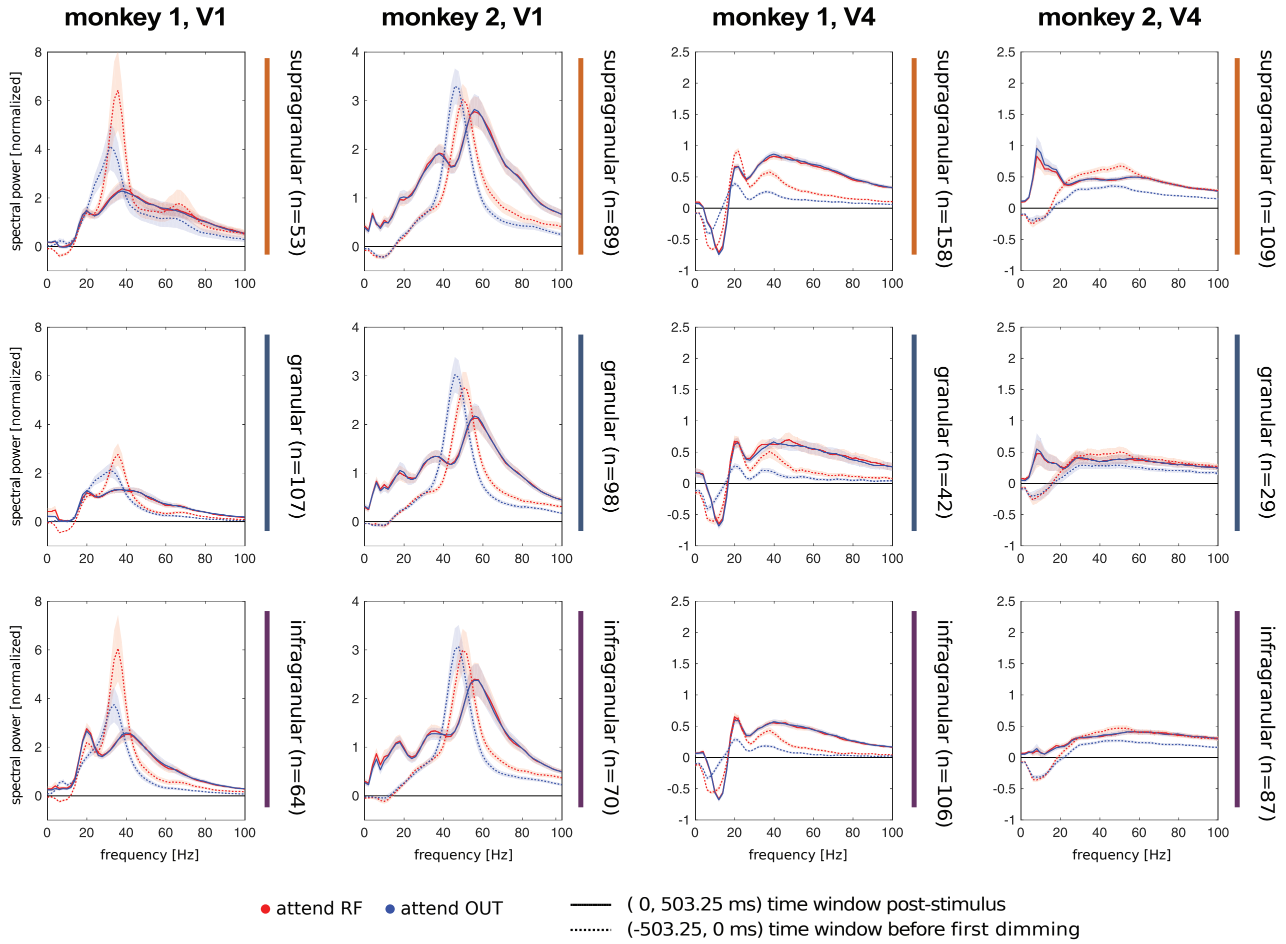


**Figure S3: Comparison of stimulus triggered LFP spectral power with attentional related spectral power.** Panels in each column show baseline-corrected spectral power (mean +/- S.E.M across sessions and depths) computed at different task-related time windows (solid lines: 0 to 503.25 ms after stimulus onset; dotted lines: -503.25 to 0 ms before first stimulus dimming), for the two attentional conditions (attend RF, attend OUT). Baseline time window is set to -203.5 to 0 ms before stimulus onset. Panels in each row refer to different laminar compartments (supragranular, granular, infragranular) for both monkeys either in V1 or in V4.

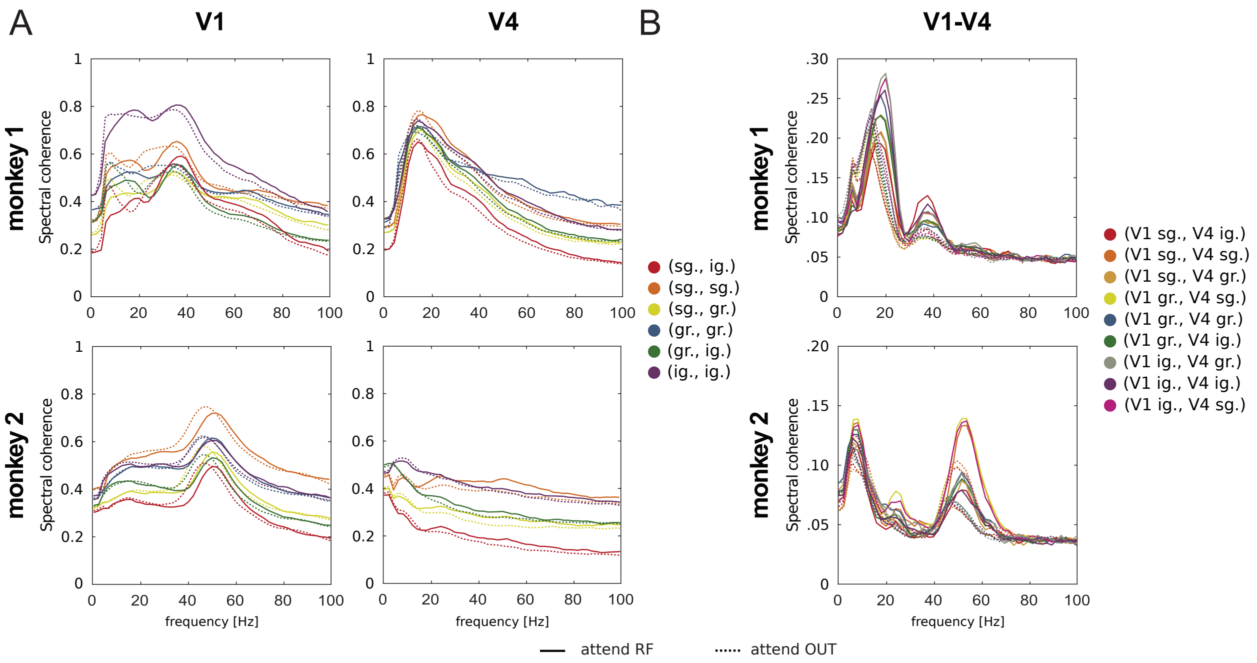


**Figure S4: LFP spectral coherence in the laminar compartments of V1 and V4 areas for the two monkeys. A)** LFP spectral coherence (mean across sessions and depths) for monkey 1 (top row) and monkey 2 (bottom row), for paired laminar depths within visual area V1 (left column) and within V4 (right column). Spectral coherence is computed on LFPs at times -503.25 to 0 ms before first stimulus dimming, and shown separately for trials with attention cued to RF location (solid lines) and for trials requiring to attend OUT (dotted lines). **B)** same as in A, but for laminar depth pairs between V1 and V4 columns. **A-B)** Laminar compartment labels are shortened: sg. (supragranular), gr. (granular) and ig. (infragranular).

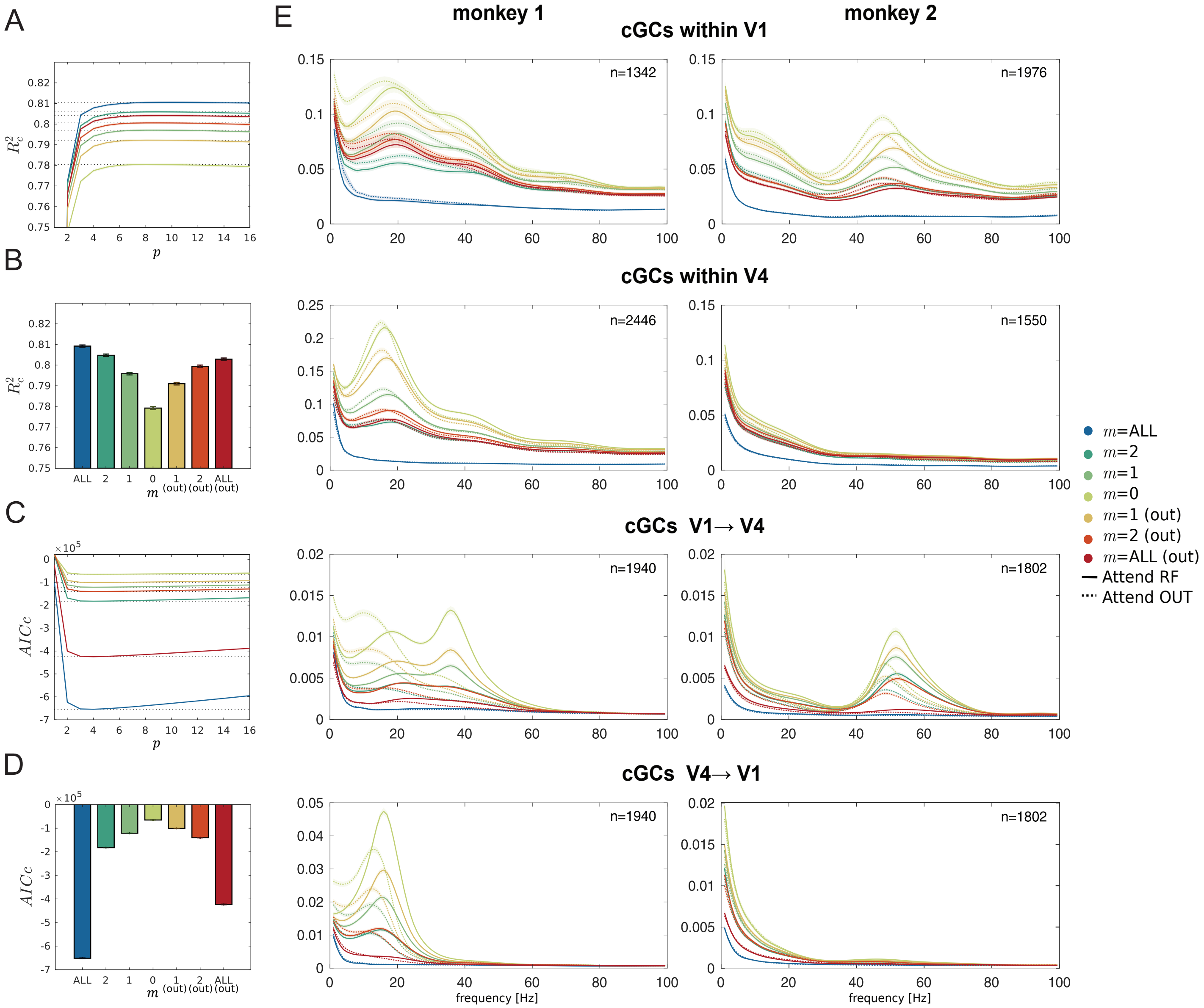


**Figure S5: Conditional GC estimation, selection of computational parameters $p$ and $m$. A)** Coefficient of determination $R_c^2$ (mean +/- S.E.M), a normalized measure of residual squared error in the estimation of VAR($p$) models, adjusted to data set size. The VAR($p$) models are estimated by least square QR decomposition for different values of VAR($p$) model order $p$ and for different conditioning strategies indexed by different values of the variable $m$. The values of $R_c^2$ are pooled for all possible V1 and V4 depth pairs (within V1, within V4, and between V1 and V4) and for the two monkeys. **B)** Average $R_c^2$ for different values of $m$ and for fixed $p$=10, yielding overall best $R_c^2$. **C)** Same as in A, but showing Akaike Information Criterion corrected for data set size ($AICc$). **D)** Same as in B, but showing $AICc$. **E)** Spectral cGCs (mean +/- S.E.M) for all possible directed pairs within V1 (first row), within V4, (second row) and between V1 and V4 (from V1 to V4 in the third row, from V4 to V1 in the fourth row), for different values of $m$, and $p$=10. For the methods tested, cGCs did not drastically change when using different values of $p$, buth their spectral resolution could increase with larger $p$. Results are reported separately for monkey 1 (left panels) and monkey 2 (right panels), and for trials where attention was cued to RF location (solid lines) and OUT (dotted lines). **A-E)** Different conditioning strategies are indexed by different values of $m$: $m$=ALL (fully conditional cGC); $m$=1, $m$=2 (respectively computed by conditioning cGC to time series in 1 or 2 most informative contacts for any directed pair); $m$=1 (out), $m$=2 (out) and $m$=ALL (out) (respectively computed by conditioning GC to time series in 1, 2 or ALL most informative contacts outside the laminar compartments of any directed pair).

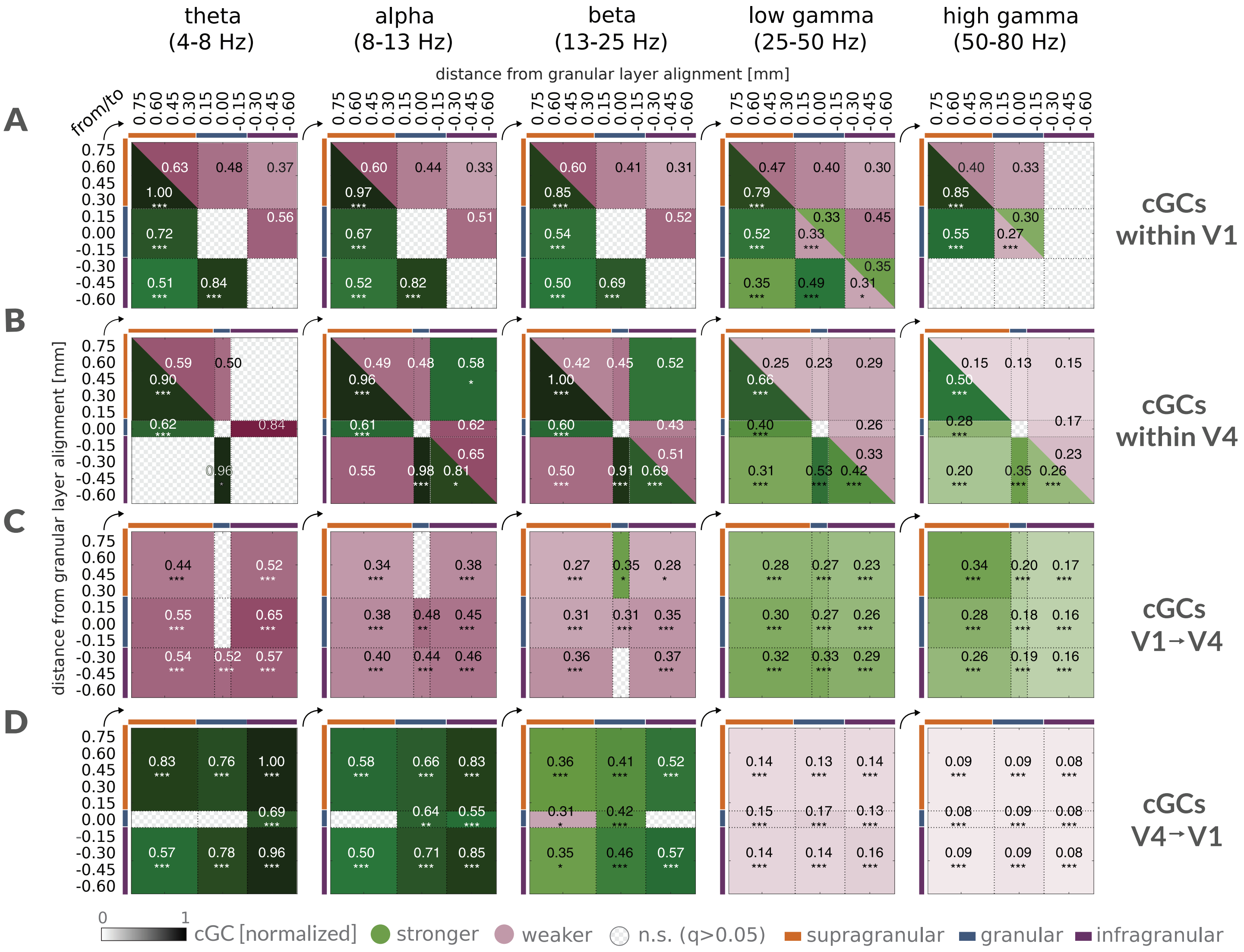


**Figure S6: Directed cGC connection matrices and dominant interactions. A)** Directed connection matrices for cGCs (mean across sessions, mean among directed contact pairs in their respective laminar compartments, pooled for the two monkeys) within V1 columns, at different frequency bands (theta: 4-8Hz, alpha 8-13 Hz, beta 13-25 Hz, low-gamma 25-50 Hz, high-gamma 50-80 Hz). Connection matrices are color coded to show significantly dominant directions (green) and non-dominant directions (magenta). The color intensity of directed connections shows the relative strength of cGCs. Significance of cGCs dominance in opposite directions is assessed by two-sided Wilcoxon signed rank tests, FDR corrected within frequency bands (* indicates q≤0.05, ** is for q≤0.01, and *** is for q≤0.001). **B)** Same as in A, but for cGCs within V4 columns. **C)** Same as in A, but for cGCs from V1 to V4 depths. **D)** Same as in A, but for cGCs from V4 to V1 depths.

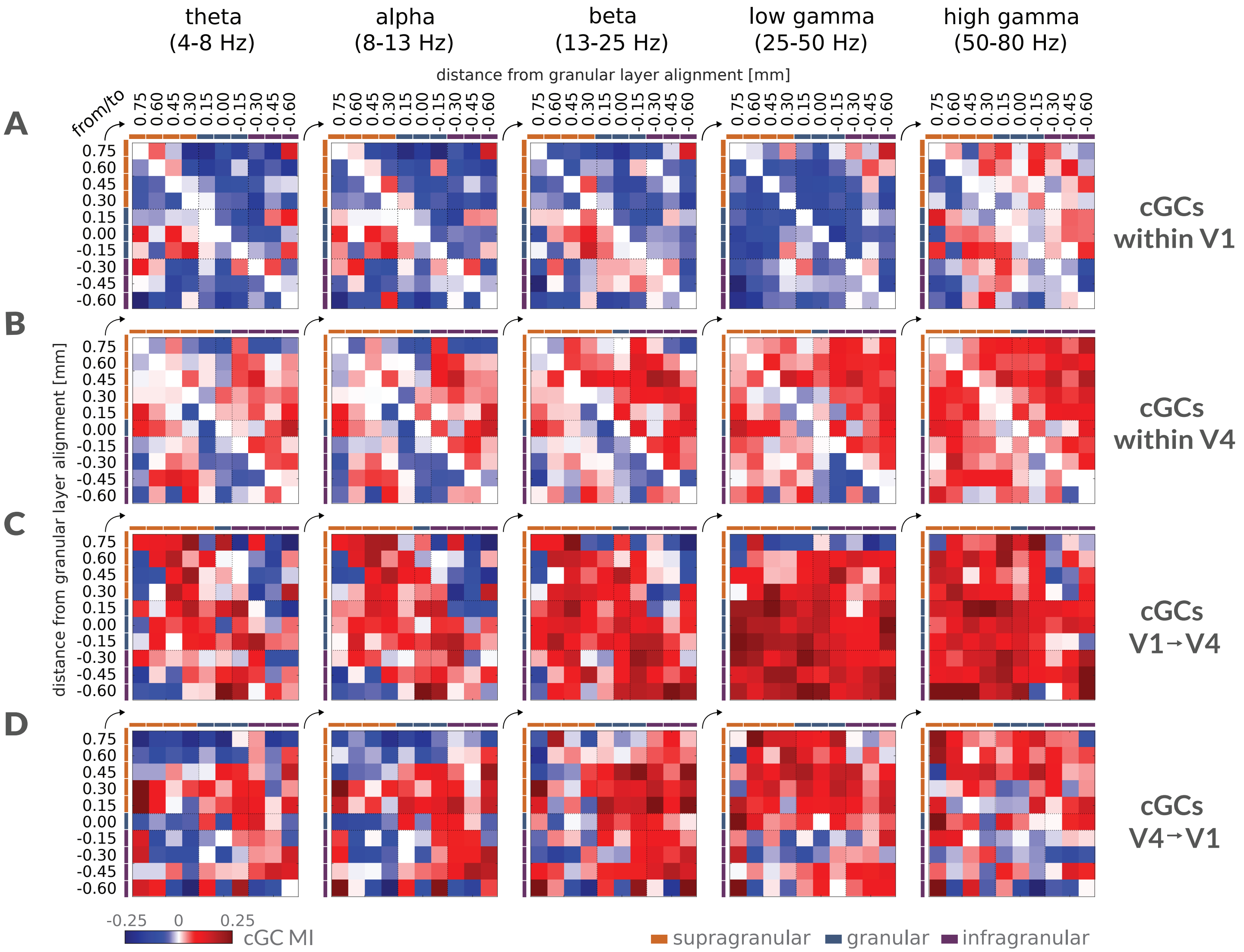


**Figure S7: Directed cGC connection matrices and attentional modulation. A)** Directed connection matrices for cGC attentional MIs (mean across sessions, pooled for the two monkeys) among directed depth pairs within V1 columns, at different frequency bands (theta: 4-8Hz, alpha 8-13 Hz, beta 13-25 Hz, low-gamma 25-50 Hz, high-gamma 50-80 Hz). **B)** Same as in A, but for cGCs within V4 columns. **C)** Same as in A, but for cGCs from V1 to V4 depths. **D)** Same as in A, but for cGCs from V4 to V1 depths.

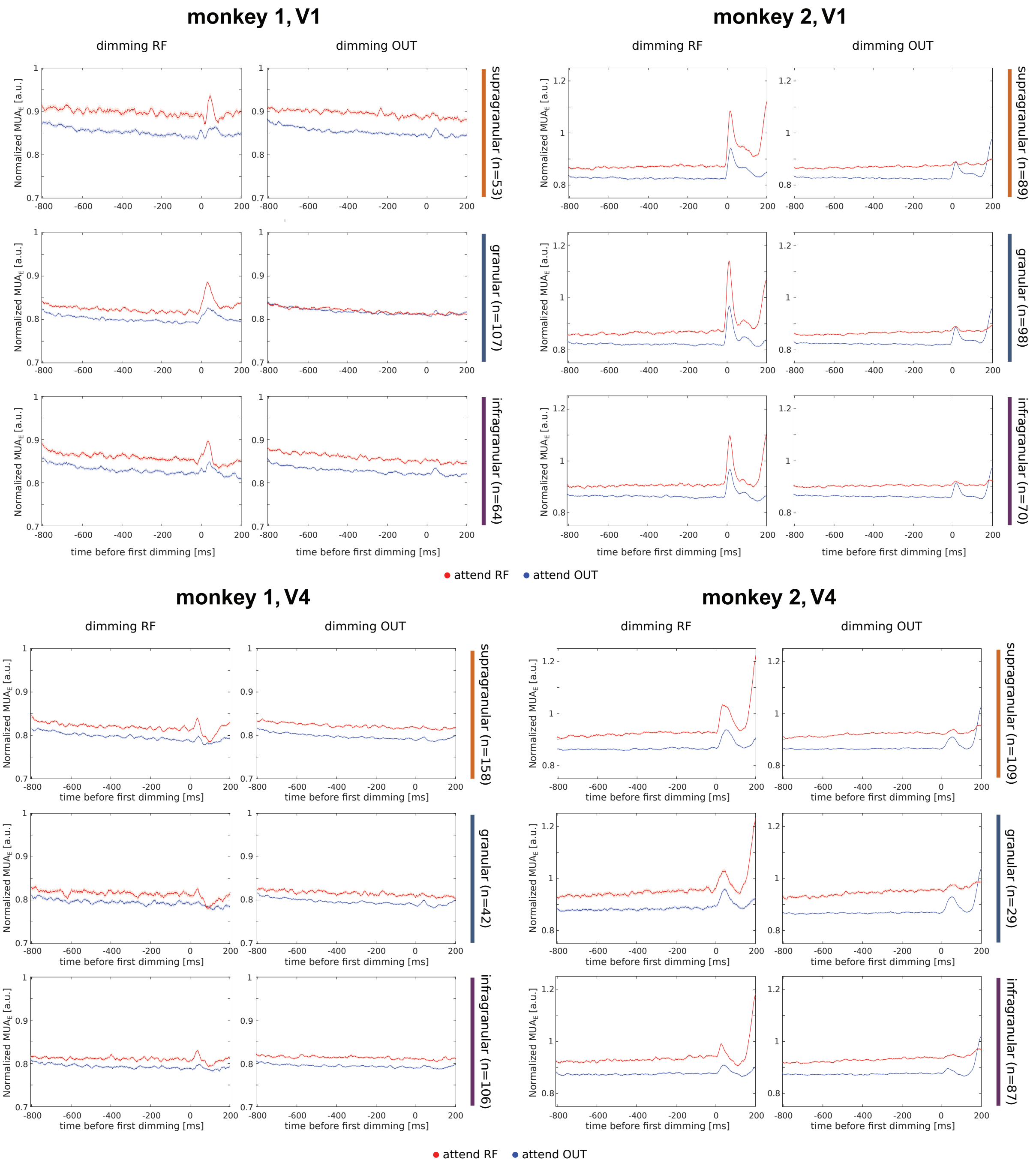


**Figure S8: Multi-unit activity envelope during attentional task behavior.** Multi-unit activity envelope ($MUA_E$; mean +/- S.E.M across sessions and depths) aligned to the timing of first stimulus dimming, and mornalized to (-200 to 0 ms) pre-stimulus activity. Results are shown separately for attentional conditions (attend RF, red lines and attend OUT, blue lines), dimming conditions (dimming RF, when first dimming occurred at RF location and dimming OUT, when first dimming occurred outside RF), and monkeys (monkey 1 two left columns, monkey 2 two right columns). Panels in different rows show results for different laminar compartments (supragranular, granular, infragranular) and for the two areas (V1 three top rows, V4 three bottom rows).
